# The MBW complex regulates volatiles in petunia flowers: EOBV interacts with AN1 to suppress biosynthesis of phenylpropenes

**DOI:** 10.64898/2026.03.01.708793

**Authors:** Oded Skaliter, Raz Cohen, Ehud Leor-Librach, Ekaterina Shor, Ori Rudich, Orit Edelbaum, Elena Shklarman, Tania Masci, Alexander Vainstein

## Abstract

Pigment production in petunia is regulated by the bHLH AN1 and the WDR protein AN11, which together with interchangeable MYBs form the MYB–bHLH–WDR (MBW) complex. Pigments and scent are interlinked flower traits, produced via the phenylpropanoid pathway. However, involvement of the MBW complex in regulating floral scent has not been demonstrated. TRV-based suppression of either *AN1* or *AN11* led to an increase in volatile emission, indicating that they are involved in negative regulation of this trait. Yeast two-hybrid and in-planta pairwise and three-way protein–protein interaction assays revealed that EMISSION OF BENZENOIDS V (EOBV) is a component of the MBW complex. Headspace and internal pool analyses of flowers from *eobv*-knockout lines, generated using a viral-based CRISPR/Cas9 system, revealed that EOBV fine-tunes volatile production: phenylpropene levels increased while those of benzenoids and phenylpropanoid-related compounds decreased. Accordingly, transcript levels of *C4H*, directing carbon flux to phenylpropenes, and *ADT3* were significantly elevated in *eobv* flowers, along with decreases in *PAAS* and *BSMT*. *EOBV* is heat-responsive and under a high-temperature regime, in addition to its involvement in scent production, it affected flower development by mitigating reduction of flower size. EOBV’s participation in the MBW complex that regulates volatiles and anthocyanins reveals an intriguing molecular link between these showy traits and flower development.

## Introduction

Petunias have evolved distinct sets of showy traits—known as pollination syndromes—to attract their specialized pollinators. These include the production of floral volatiles, such as those emitted by *Petunia axillaris* to attract moths, and pigmentation, with purple flowers in *Petunia inflata* and red ones in *Petunia exserta*, which attract bees and hummingbirds, respectively (Gübitz, T. Hoballah, M.E. Dell’Olivo, A. Kuhlemeier, 2009). Both volatiles and pigments are primarily synthesized from the aromatic amino acid phenylalanine via the phenylpropanoid pathway (Skaliter et al., 2022) through a complex and interlinked network of enzymes and regulators (Fig. 1). The extensive crosstalk between these metabolic branches, yielding phenylalanine-derived volatiles and anthocyanin pigments, has been demonstrated in numerous studies (Zuker et al., 2002; Zvi et al., 2008; Zvi et al., 2012; Cna’ani et al., 2015; Shaipulah et al., 2016; Ravid et al., 2017; Patrick et al., 2021).

**Fig. 1.**
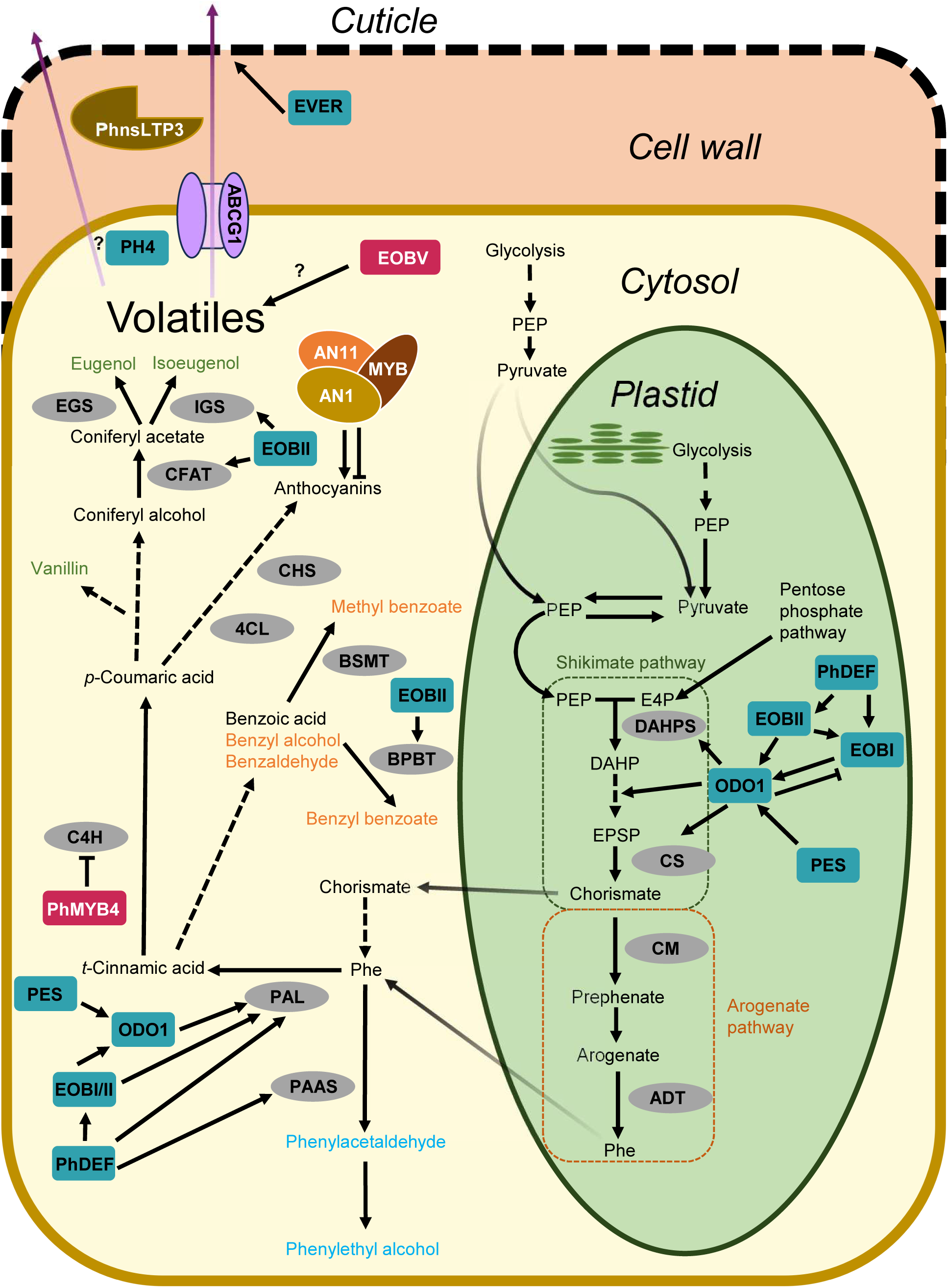
Overview of phenylpropanoid biosynthetic pathways leading to volatile production and emission in petunia flowers. L-phenylalanine (Phe) is synthesized in the plastid or in the cytosol from chorismate generated via the shikimate pathway and arogenate pathways from. From Phe, distinct enzymatic reactions lead to the production of volatiles: benzenoids (C6–C1), shown in orange; phenylpropanoid-related compounds (C6–C2), shown in teal; and phenylpropenes (C6–C3), shown in green. Following their biosynthesis, volatiles are emitted (indicated by purple arrows) across the plasma membrane via the ATP-binding cassette (ABC) transporter PhABCG1 and through the cell wall via the non-specific lipid-transfer protein 3 (PhnsLTP3), before crossing the cuticle. The EPIDERMIS VOLATILE EMISSION REGULATOR (EVER) has been shown to control volatile emission by modulating wax content through the regulation of epicuticular wax biosynthesis. In addition, PH4 has also been shown to affect volatile emission through a yet unknown mechanism. Alternatively, from *p*-Coumaric acid, multiple enzymatic steps lead to the production of anthocyanins, whose biosynthesis is regulated by the MYB–bHLH–WDR (MBW) complex consisting of the bHLH AN1, the WDR protein AN11, and interchangeable MYBs. Enzymes are shown in gray circles; positive and negative transcriptional regulators are shown in teal and red rectangles, respectively. Stacked arrows represent multiple enzymatic steps. Abbreviations: arogenate dehydratase (ADT); benzoyl-CoA:benzylalcohol/2-phenylethanol benzoyltransferase (BPBT); benzoic acid/salicylic acid carboxyl methyltransferase (BSMT); chalcone synthase (CHS); chorismate mutase (CM); chorismate synthase (CS); cinnamate 4-hydroxylase (C4H); coniferyl alcohol acetyltransferase (CFAT); 4-coumarate-CoA ligase (4CL); deficiens (PhDEF); 3-deoxy-D-arabino-heptulosonate-7-phosphate (DAHP); 3-deoxy-D-arabino-heptulosonate-7-phosphate synthase (DAHPS); emission of benzenoids I/II/V (EOBI/II/V); 5-3nolpyruvylshikimate-3-phosphate (EPSP); erythrose-4-phosphate (E4P); eugenol synthase (EGS); isoeugenol synthase (IGS); odorant 1 (ODO1); phenylacetaldehyde synthase (PAAS); L-phenylalanine ammonia-lyase (PAL); phenylpropanoid emission-regulating scarecrow-like (PES); phosphoenolpyruvate (PEP).

The scent bouquet of petunia is comprised of volatiles methyl benzoate, benzaldehyde, benzyl benzoate and benzyl alcohol, which are produced via the C6-C1 (benzenoid) branch following the action of L-PHENYLALANINE AMMONIA LYASE (PAL). In addition, petunia flowers emit C6-C2 volatiles (phenylpropanoid-related compounds) such as phenylethyl alcohol and phenylacetaldehyde, and phenylpropenes (C6–C3) such as eugenol, isoeugenol and vanillin (Fig. 1) (Muhlemann et al., 2014). Production and emission of volatiles are tightly regulated at multiple levels: spatial regulation (Skaliter et al., 2021), physical cellular barriers (Liao et al., 2021; Liao et al., 2023; Skaliter et al., 2024), environmental cues such as ambient temperature and light (Cna’ani et al., 2014; Shor et al., 2023a) and cellular compartmentalization (Cna’ani et al., 2017). Furthermore, involvement of the phytohormones ethylene, gibberellin (GA), auxin and jasmonic acid in the regulation of scent production and emission has been described (Underwood et al., 2005; Liu et al., 2017; Ravid et al., 2017; Lynch et al., 2020; Skaliter et al., 2024). The molecular factors underlying these levels of regulation belong to different families: adenosine triphosphate–binding cassette (ABC) transporters (Adebesin et al., 2017), GRAS (Ravid et al., 2017; Shor et al., 2023a), non-specific lipid transfer proteins (Liao et al., 2023), ethylene-response factors (Liu et al., 2017), basic helix–loop–helix (bHLH) (Shor and Vainstein, 2024) and MADS-box (Bednarczyk et al., 2025). The dominant group of transcriptional regulators of floral volatiles belongs to the MYB superfamily (Verdonk et al., 2005; Spitzer-Rimon et al., 2010; Colquhoun et al., 2011; Spitzer-Rimon et al., 2012; Cna’ani et al., 2015; Skaliter et al., 2024). These regulators bind DNA through their highly conserved N-terminal MYB domain and, depending on their C-terminal domain, can activate or repress target genes (Dubos et al., 2010). EMISSION OF BENZENOIDS I and II (EOBI and EOBII, respectively) and ODORANT1 (ODO1) promote the expression of genes encoding enzymes responsible for phenylalanine and dedicated volatiles’ biosynthesis, whereas PH4 is required specifically for the volatiles’ emission (Verdonk et al., 2005; Spitzer-Rimon et al., 2010; Spitzer-Rimon et al., 2012; Cna’ani et al., 2015). Another MYB activator, EPIDERMIS VOLATILE EMISSION REGULATOR (EVER), is involved in repressing low-vapor-pressure volatiles’ emission by modulating the composition of petal epicuticular waxes (Skaliter et al., 2024). The repressor PhMYB4 downregulates transcripts of *CINNAMATE 4-HYDROXYLASE* (*C4H*), the enzyme that channels carbon flux to phenylpropenes (Colquhoun et al., 2011). Based on general screening of floral MYBs using virus-induced gene silencing (VIGS) in petunia cv. P720, EOBV has also been suggested as a repressor of volatiles (Spitzer-Rimon et al., 2012), but its mode of action and targets are unknown (Fig. 1).

Similar to volatiles, the biosynthesis of anthocyanins is regulated by an intricate network of MYBs. To exert their action, these MYBs interact with bHLH and WD repeat-containing proteins (WDR, also known as WD40) to form the MBW complex (Quattrocchio et al., 1999; Albert et al., 2011; Albert et al., 2014; Zhang et al., 2019). This complex’s functionality has been shown to require all members, as mutations in any one of them results in acyanic petals (de Vetten et al., 1997; Quattrocchio et al., 1999; Spelt et al., 2000). In this complex, MYBs provide target-gene specificity, bHLH proteins function as transcriptional co-activators and serve as a physical bridge between MYB and WDR components, and WDR stabilizes the complex and promotes its assembly (Xu et al., 2015; Bulanov et al., 2025). In petunia, the complex typically consists of the bHLH ANTHOCYANIN1 (AN1) and the WDR ANTHOCYANIN11 (AN11), whereas the MYB component that dictates the target and activity of the complex depends on the tissue, environmental conditions or both (Albert et al., 2011; Albert et al., 2014; Zhang et al., 2019). While involvement of the MBW complex in the regulation of phenylpropanoid-derived volatiles’ production has not been demonstrated, UNIQUE PLANT PHENYLPROPANOID REGULATOR, a Viridiplantae-specific protein containing a combination of WDR, kinase and RING domains, was recently shown to regulate this process in petunia flowers (Shor et al., 2023b). Moreover, in seeds of sorghum (*Sorghum bicolor* L. Moench), a homologous MBW complex was shown to regulate both anthocyanins and fatty acid-derived volatiles (Xie et al., 2019).

Here we show involvement of the petunia MBW complex in the regulation of floral volatiles. Using AN1 as bait, we reveal that EOBV is a component of this complex; *eobv*-knockout flowers accumulated higher levels of C6-C3 and lower levels of C6-C1/C2 compounds mainly by controlling *C4H* expression. Under high ambient temperatures, flower size and volatile production in eobv-knockout lines were significantly less affected compared to controls, suggesting that EOBV integrates environmental and developmental cues. These results unravel EOBV’s mode of action, and by showing its participation in the MBW complex, further demonstrate the interconnection of the regulatory mechanisms underlying the biosynthesis of volatiles and anthocyanins in flowers.

## Results

### VIGS-mediated suppression of *AN1* and *AN11* enhances floral volatile emission in petunia

Previous studies have shown that core components of the MBW complex, *AN1* and *AN11*, are expressed in young flowers (de Vetten et al., 1997; Spelt et al., 2000). To detail their developmental expression pattern in later stages of flower development, when the volatile machinery is active, RNA was extracted from petal tissues ranging from 4-cm buds to 4 days postanthesis (DPA) and analyzed by reverse transcription quantitative PCR (RT-qPCR). Similar to the early stages, *AN1* and *AN11* showed overlapping expression patterns, which aligned with volatile production and emission—peaking at anthesis, remaining high at 1DPA and 2DPA, and declining at 3DPA and 4DPA (Fig. 2a). To evaluate the involvement of the MBW complex in volatile production in petunia, *AN1* and *AN11* were individually suppressed in cv. Mitchell flowers. To achieve this, a 224-bp fragment of *AN1*’s 5’ untranslated region (UTR), and 332 bp of the 3’ UTR of *AN11*, were amplified from cDNA generated from flowers and cloned into a pTRV2 vector to transiently downregulate their expression using VIGS (Figs. 1a and 1b). Flowers at anthesis were inoculated with *Agrobacterium* carrying pTRV2-AN1, pTRV2-AN11 or pTRV2-CHALCONE SYNTHASE (CHS) as a control (Spitzer et al., 2007), followed by localized headspace sampling (Skaliter et al., 2021). GC–MS analysis revealed a significant increase in emission of the phenylpropenes eugenol and vanillin, the C6-C1 branch compounds benzyl alcohol and methyl benzoate as well as the C6-C2 compound phenylethyl alcohol, in petal tissues inoculated with pTRV2-AN1 and pTRV2-AN11 as compared to control pTRV2-CHS (Fig. 2b). In pTRV2-AN11, emission of (*Z*)-isoeugenol was also enhanced. RT-qPCR analysis (Fig. 2c) and VIGS suppression in anthocyanin-accumulating cv. P720 flowers (Fig. S1c) confirmed the suppression activity of the *AN1* and *AN11* constructs. Taken together, these results indicated that while AN1 and AN11 are necessary for anthocyanin biosynthesis, they also negatively affect volatile production.

**Fig. 2.**
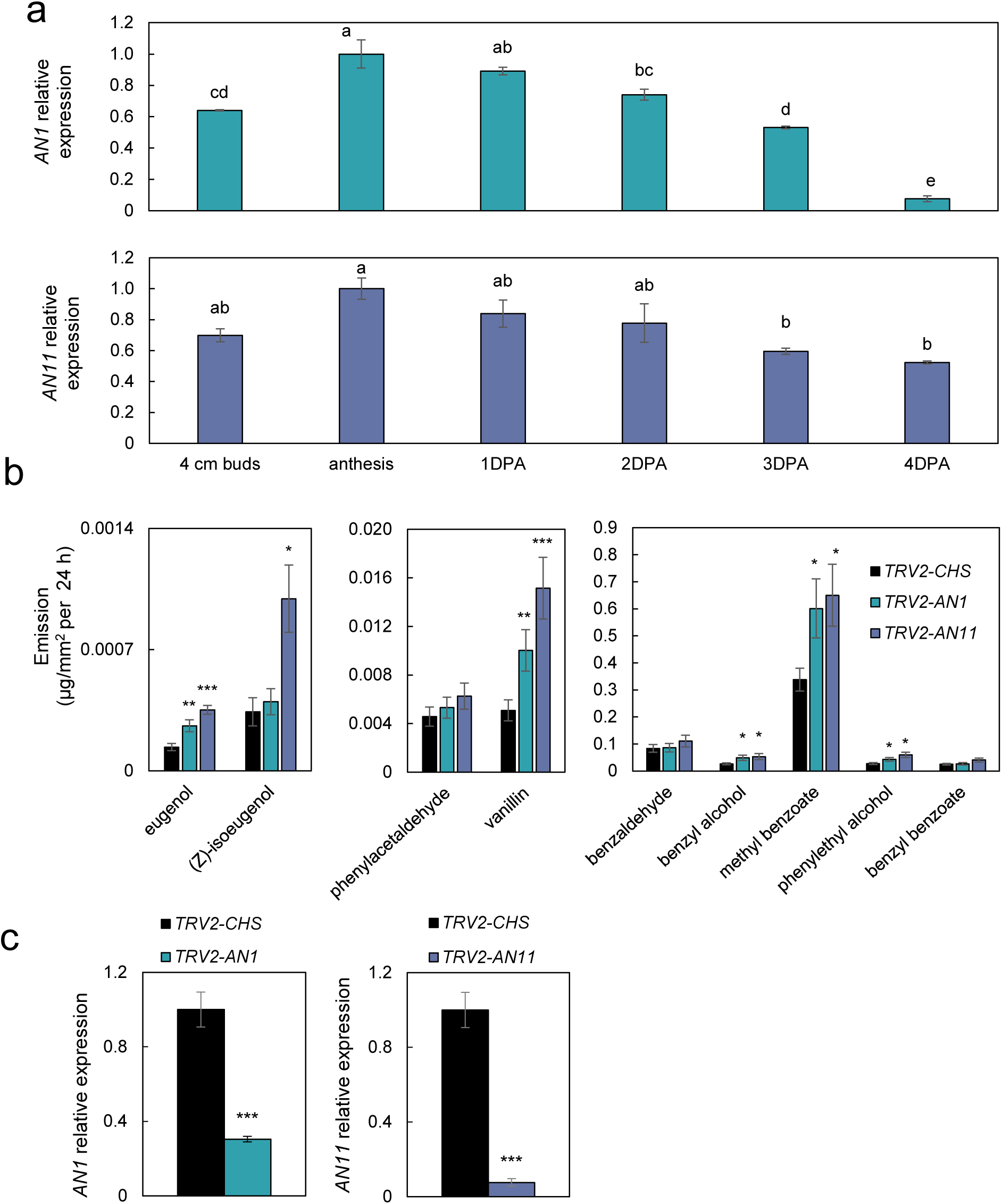
Petunia MBW complex components AN1 and AN11 are involved in negative regulation of volatiles. **a)** Developmental expression patterns of *AN1* and *AN11*. RT-qPCR analysis was performed on flower tissues collected at different developmental stages. DPA, days postanthesis. Data are means ± SEM (n = 3). Significance of differences was calculated by Tukey’s multiple comparison test following one-way ANOVA. Values with different letters are significantly different at *P* ≤ 0.05. **(b and c)** Petunia cv. Mitchell flowers were inoculated at anthesis with agrobacteria carrying pTRV2-AN1, pTRV2-AN11 or pTRV2-CHS as a control. Flowers were harvested 2 days after inoculation and subjected to localized headspace sampling followed by GC–MS analysis **(b)** or RNA extraction followed by RT-qPCR analysis **(c)**. Data are means ± SEM (*n* = 12). Significance of differences was calculated using Dunnett’s test with pTRV2-CHS as the control following one-way ANOVA **(b)** or two-tailed unpaired Student’s *t*-test **(c)** (\**P* ≤ 0.05, \*\**P* ≤ 0.01, \*\*\**P* ≤ 0.001). **(a and c)** RT-qPCR data were normalized to *ACTIN*, and the presented data were normalized to the sample with the highest expression level. Standard errors are indicated by vertical lines.

### AN1 interacts with the MYB volatile regulator EOBV

To identify MYB volatile regulators that may interact with AN1 and AN11, yeast two-hybrid (Y2H) assays were performed. The MYB activators EOBI, EOBII, ODO1 and EVER, and repressors PhMYB4 and EOBV, were tested for interactions. The well-documented interaction between AN1 and AN11 was used as a positive control (Albert et al., 2014). As MYB activators can self-activate the reporter gene *LacZ* when fused to the GAL4 DNA-binding domain (BD), they were fused to the GAL4 activation domain (AD). None of the tested activators interacted with either AN1 or AN11 (Fig. S2). Similarly, there was no interaction between AN1 or AN11 and PhMYB4. In contrast, activation of the target *LacZ* in yeast transformed with EOBV fused to either AD or BD indicated that it interacts with AN1 (Fig. 3a). In line with these results, the signature R/B-like bHLH motif [D/E]Lx2[R/K]x3Lx6Lx3R, which is responsible for interactions of MYBs with bHLH, was identified in the protein sequence of EOBV (Fig. S3) (Zimmermann et al., 2004).

**Fig. 3.**
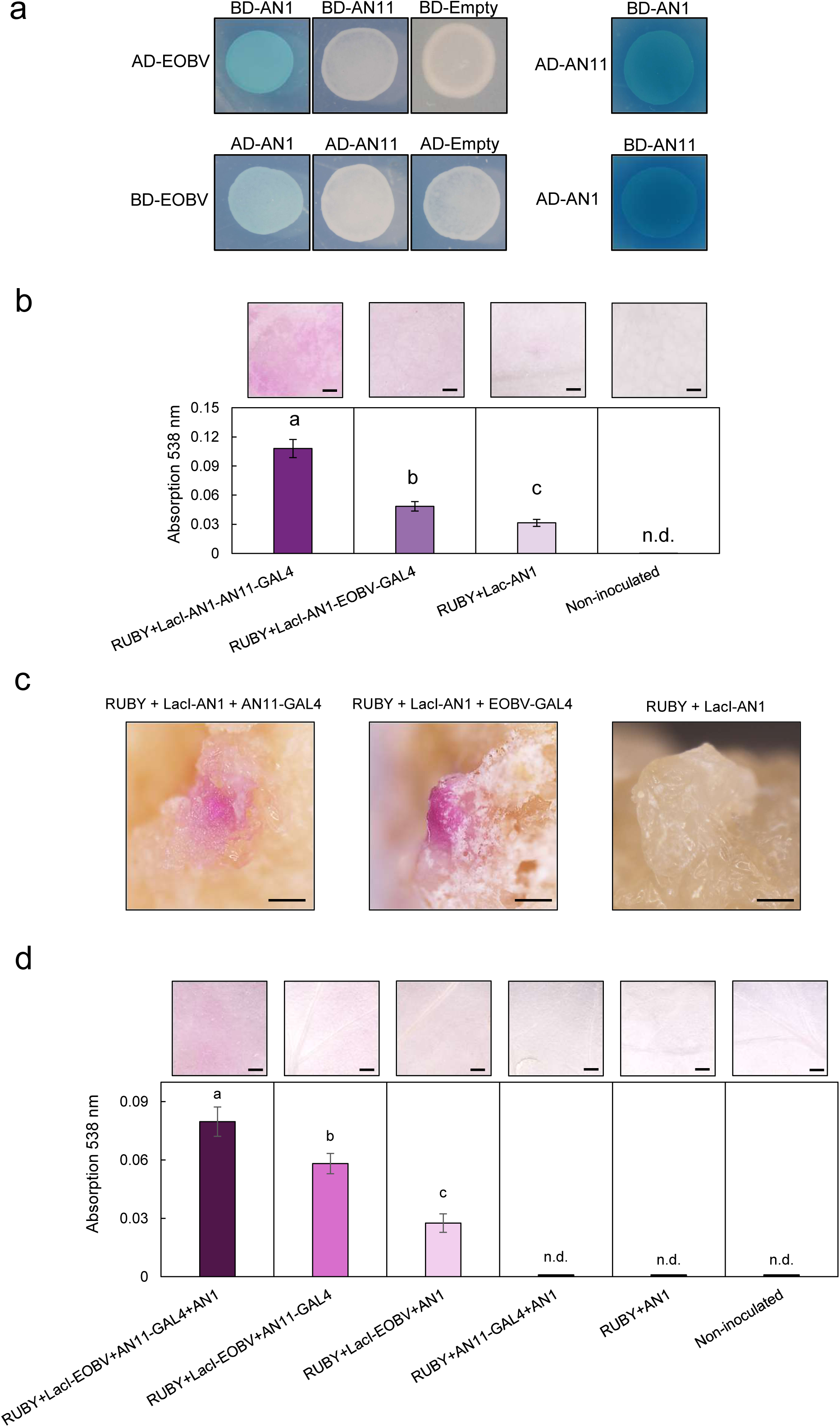
The MYB EOBV is a component of the MBW complex. **a)** Yeast-two hybrid assay was performed by fusing the coding sequences of target proteins to GAL4 binding domain (BD) or GAL4 activation domain (AD), followed by yeast transformation. X-Gal staining was performed on transformed yeast to detect *β*-galactosidase activity, indicating an interaction between bait and prey. The interaction between AN1 and AN11 was used as a positive control (Albert et al., 2014). Vectors without coding sequences (AD:empty, BD:empty) were used as negative controls. **(b–c)** In-planta pairwise protein-interaction assays. **b)** *N. benthamiana* leaves were inoculated with a mixture of agrobacteria carrying the vectors shown in Supplementary Figure 4a; 5 days postinoculation, leaves were harvested and analyzed following immersion in 100% ethanol. Upper panel: representative photos of bleached leaves inoculated with OpLacI:RUBY, 35S:LacI-AN1 and 35S:AN11-GAL4 (positive control); OpLacI:RUBY, 35S:LacI-AN1 and 35S:EOBV-GAL4; OpLacI:RUBY and 35S:LacI-AN1 (negative control); and non-inoculated leaf. Bar = 1 mm. Lower panel: betalain levels in leaves inoculated with the above-described vectors **(c)** Left to right: representative photos of pigmented calli formed on *N*. *tabacum* leaf disks inoculated with OpLacI:RUBY, 35S:LacI-AN1 and 35S:AN11-GAL4 (positive control); OpLacI:RUBY, 35S:LacI-AN1 and 35S:EOBV-GAL4; OpLacI:RUBY and 35S:LacI-AN1 (negative control). Bar = 0.5 mm. **d)** Three-way in-planta interaction assay. *N. benthamiana* leaves were inoculated with a mixture of agrobacteria carrying the vectors shown in Supplementary Figure 4A and analyzed as described in **(b)**. Upper panel: representative photos of bleached leaves inoculated with OpLacI:RUBY, 35S:LacI-EOBV, 35S:AN11-GAL4 and 35S:AN1; OpLacI:RUBY, 35S:LacI-EOBV and 35S:AN11-GAL4; OpLacI:RUBY, 35S:LacI-EOBV and 35S:AN1; OpLacI:RUBY, 35S:AN11-GAL4 and 35S:AN1; OpLacI:RUBY and 35S:AN1; and non-inoculated leaf. Bar = 1 mm. Lower panel: betalain levels in leaves inoculated with the above-described vectors. Data are means ± SEM (*n* = 5). Significance of differences was calculated by Tukey’s multiple comparison test following one-way ANOVA. Values with different letters are significantly different at *P* ≤ 0.05; n.d., not detected. Standard errors are indicated by vertical lines.

Similar to studies on several other MYBs, EOBV did not interact with AN11 when fused to AD or BD (Albert et al., 2014). To further confirm the interaction between EOBV and AN1, in-planta protein–protein interaction assays, using *RUBY* as the reporter gene, were conducted (He et al., 2020; Chen et al., 2023). The assay was based on Chen et al. (2023), except that *RUBY* was placed under the control of six repeats of the lac operator fused to a minimal 35S promoter (6×OpLacI:mini35S), whereas the bait (AN1) was fused to LacI BD, and EOBV and AN11 were each fused to GAL4 AD, all driven by the cauliflower mosaic virus (CaMV) 35S promoter (Fig. S4a). Leaves of *Nicotiana benthamiana* were inoculated with *Agrobacterium* carrying 6×OpLacI:mini35S:*RUBY* together with LacI-AN1 and EOBV-GAL4, or LacI-AN1 and AN11-GAL4 as a positive control, or only LacI-AN1 as a negative control. Betalains were observed in the inoculated tissues 5 days postinoculation, and were extracted and quantified. The analysis revealed accumulation of betalains in LacI-AN1/EOBV-GAL4-inoculated leaves, indicating interaction between the proteins (Fig. 3b). In LacI-AN1/AN11-GAL4-inoculated tissues, betalains accumulated to higher levels than in the LacI-AN1/EOBV-GAL4-inoculated ones, suggesting a stronger interaction. These results are in line with the Y2H assay, which also revealed a stronger interaction between AN1 and AN11 than between AN1 and EOBV. Leaves inoculated only with LacI-AN1 (negative control) showed significantly lower betalain accumulation. To further confirm the interaction between AN1 and EOBV, leaf disks (30 per treatment) of *Nicotiana tabacum* were inoculated with *Agrobacterium* carrying the above constructs and placed on regeneration media. After 2 weeks, inoculated leaf disks were bleached in ethanol and the development of betalain-accumulating calli on the disks was monitored. Multiple pigmented calli were observed on leaf disks inoculated with LacI-AN1/EOBV-GAL4, as well as on those inoculated with LacI-AN1/AN11-GAL4 (Fig. 3c). No pigmented calli were observed on LacI-AN1-inoculated disks. Taken together, the Y2H and in-planta interaction assays demonstrated that EOBV interacts with AN1 of the MBW complex.

To test whether EOBV forms a ternary complex with AN1 and AN11, we devised a three-way in-planta protein–protein interaction assay (Fig. S4a). Given the lack of direct interaction between EOBV and AN11 in pairwise assays, we examined whether they can activate 6×OpLacI:mini35S:*RUBY* reporter in the presence of AN1, acting as a bridge to facilitate MBW complex formation. Two new plasmids were generated: p35S:AN1 with pUBQ:DsRed (as a reporter) and p35S:LacI-EOBV (Fig. S4a). *N*. *benthamiana* leaves were infiltrated with different construct combinations to evaluate the in-planta formation of an active EOBV–AN1–AN11 complex. Co-infiltration of all three components—LacI-EOBV, AN11-GAL4, and AN1—resulted in prominent *RUBY* reporter activation, leading to visible betalain accumulation 48 h postinfiltration (Fig. S4b). In line with these results, betalain extraction and quantification showed that tissues inoculated with LacI-EOBV/AN11-GAL4/AN1 accumulated the highest levels of betalains compared to all other treatments (Fig. 2d). Inoculation of leaves with only two components, LacI-EOBV/AN11-GAL4 or LacI-EOBV/AN1, resulted in significantly lower betalain accumulation, whereas no betalains were detected in tissues inoculated with AN11-GAL4/AN1 or AN1 alone. DsRed expression was observed in all AN1-containing treatments (Fig. S4b). To further confirm the ability of the triad to activate *RUBY*, *N*. *tabacum* disks were inoculated with LacI-EOBV, AN11-GAL4 and AN1 constructs. Pigmented regions were observed in inoculated explants after 2 weeks, demonstrating their ability to activate *RUBY* by forming a complex (Fig.4c). Taken together, these results indicated that EOBV participates in the MBW complex with AN1 and AN11.

### EOBV is a petal-specific repressor that belongs to MYB family subgroup 4

To characterize the evolutionary relationship of EOBV in the MYB family, we performed a phylogenetic analysis of MYBs from various plant species. Phylogenetic analysis clustered EOBV into MYB family subgroup 4 (Dubos et al., 2010), which contains previously characterized MYB repressors (Figs. 4a and S5) (Ma and Constabel, 2019). More specifically, EOBV clustered in the SG4-AtMYB subclass together with MYBs regulating phenylpropanoid biosynthesis (Jin et al., 2000; Colquhoun et al., 2011; Zhang et al., 2023). Its closest homolog was tomato (*Solanum lycopersicum*) MYB7, shown to be involved in negative regulation of anthocyanins (Zhang et al., 2023). InterPro analysis (Blum et al., 2021) of EOBV revealed that similar to most subgroup 4 members, EOBV has two conserved motifs at its C terminus: C1 (LlsrGIDPxT/sHRxI/L) and C2 (pdLNLD/ELxiG/S), the latter part of the ethylene-responsive element binding factor-associated amphiphilic repression (EAR) motif which is responsible for the repression activity (Fig. S3) (Ohta et al., 2001; Chen et al., 2019). EOBV also has a zinc-finger domain C3 motif (CX_1–2_CX_7–12_CX_2_C) and a C4 motif (FLGLx4-7V/LLD/GF/YR/Sx1LEMK), which is typical for the SG4-AtMYB subclass (Fig. S3) (Chen et al., 2019). To determine whether, like most MYBs, EOBV is located in the nucleus, it was fused to green fluorescent protein (GFP) at either the N or C terminus and transiently expressed in petunia petals. Confocal microscopy confirmed that the GFP signal of EOBV-GFP and GFP-EOBV co-localizes with a red fluorescent protein (RFP) fused to a nuclear localization signal (NLS), which served as a positive control for nuclear localization (Fig. 4b).

**Fig. 4.**
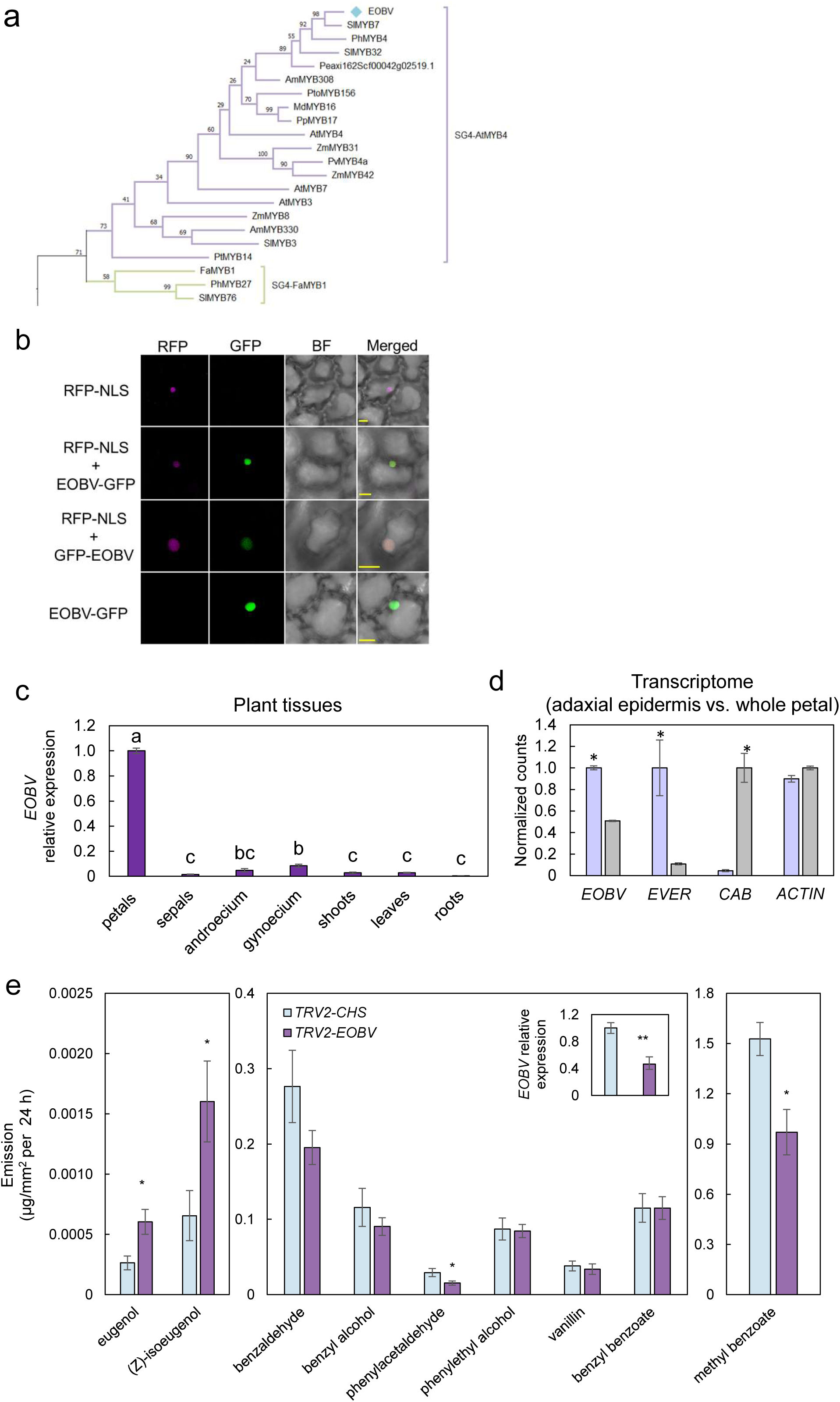
EOBV is a petal epidermis-enriched R2R3-MYB subgroup 4 member that negatively regulates volatiles in petunia flowers. **a)** Phylogenetic tree displaying the similarity between EOBV and subgroup 4 R2R3-MYBs from other plants. Protein sequences were aligned by ClustalW, followed by neighbor-joining method with 1000 bootstraps using MEGA version 11. **b)** Nuclear localization of EOBV in petal epidermal cells by confocal microscopy. Petunia petals were inoculated with agrobacteria carrying *EOBV-GFP* or *GFP-EOBV* and *RFP*-*NLS,* or only *RFP*-*NLS* as a control. Bars = 10 μm. BF = bright field. NLS, nuclear localization signal.**c)** Expression of *EOBV* in different plant organs. Tissues from different organs were collected at 1100 h. RT-qPCR data were normalized to *ACTIN*, and the presented data were normalized to the sample with the highest expression level. Significance of differences was calculated by Tukey’s multiple comparison test following one-way ANOVA. Values with different letters are significantly different at *P* ≤ 0.05. **d)** *EOBV* transcripts are enriched in the petal adaxial epidermis. Normalized counts in the petal adaxial epidermis (purple bars) vs. whole petal (gray bars) based on the transcriptome data from Skaliter *et al*. (2024). Significance of differences was calculated by two-tailed unpaired Student’s *t*-test: \**P* ≤ 0.05. EVER, EPIDERMIS VOLATILE EMISSION REGULATOR. CAB, CHLOROPHYLL A/B BINDING PROTEIN. **e)** Transient suppression of *EOBV* in petunia cv. Mitchell flowers inoculated with agrobacteria carrying pTRV2-CHS as a control or pTRV2-EOBV. Flowers were harvested 2 days after inoculation and subjected to localized headspace sampling followed by GC–MS analysis or RNA extraction followed by RT-qPCR analysis (inset). Data are means ± SEM (*n* = 11–12). RT-qPCR data were normalized to *ACTIN*, and the presented data were normalized to those from pTRV2-CHS. Significance of differences between treatments was calculated by two-tailed unpaired Student’s *t*-test: (\**P* ≤ 0.05, \*\**P* ≤ 0.01, \*\*\**P* ≤ 0.001). Standard errors are indicated by vertical lines.

To detail the spatial expression profile of *EOBV*, RT-qPCR was performed on RNA extracted from various vegetative and floral tissues. *EOBV*’s expression was found to be petal-specific, with transcript levels in the other tissues—androecium, gynoecium, sepals, roots, shoots and leaves—at substantially lower or background levels (Fig. 4c). Furthermore, analysis of the petunia petal adaxial epidermis transcriptome (Skaliter et al., 2024)—the main site of volatile emission (Skaliter et al., 2021)—revealed enrichment of *EOBV* transcript, with approximately 1.8-fold higher expression than in the whole petal (Fig. 4d). Analysis of the petal transcriptome (Shor et al., 2023b) from petunia buds and 1DPA flowers, in the morning vs. evening, revealed highest *EOBV* expression in flowers during the evening (Fig. S6). Its diurnal pattern changed according to developmental stage: higher in the morning vs. evening in buds and vice versa in 1DPA flowers.

### TRV-based suppression of *EOBV* in petunia flowers alters volatile composition

To evaluate EOBV’s effect on volatiles, a 164-bp fragment of its 3’ UTR was amplified and cloned into pTRV2 (Fig. S7a). ‘Mitchell’ flowers at anthesis were inoculated with *Agrobacterium* carrying either TRV2-EOBV or TRV2-CHS. Localized headspace analysis revealed a significant increase in the phenylpropenes eugenol and (*Z*)-isoeugenol, with a concomitant decrease in emission of methyl benzoate and phenylacetaldehyde in TRV2-EOBV-inoculated tissues (Fig. 4e). RT-qPCR analysis confirmed that *EOBV* transcripts were reduced by ca. 60% in these tissues compared to control TRV2-CHS. To test whether EOBV is involved in anthocyanin regulation, plants of petunia cv. Classic Blue Ray, producing purple-pigmented flowers, were inoculated with the aforementioned TRV2 vectors. Extraction of total anthocyanins from flowers of plants inoculated with TRV2-EOBV revealed that, in contrast to its homolog in tomato, suppression of *EOBV* does not affect anthocyanin content (Fig. S7b). GA has been shown to repress volatile and induce anthocyanin production in petunia flowers (Ravid et al., 2017). To determine GA’s effect on *EOBV* expression, 4-cm buds were treated with exogenous GA₃, followed by RT-qPCR analysis. Transcript levels of *GIBBERELLIN-INDUCED PROTEIN 2*, used as a positive control for GA application, increased following GA treatment (Fig. S8) (Ben-Nissan et al., 2004; Skaliter et al., 2024). In line with the observed lack of EOBV effect on anthocyanin levels, its transcript level was unaffected by GA₃ application, suggesting that *EOBV* is not regulated by GA.

### EOBV controls phenylpropene production through transcriptional suppression of *C4H*

To detail EOBV’s involvement in the regulation of volatiles production and emission, knockout mutations were induced in petunia cv. Mitchell using the viral-based CRISPR/Cas9 system (Skaliter et al., 2024). The petunia genome has a single copy of *EOBV* that is comprised of two exons and one intron (Fig. 5a) (Bombarely et al., 2016). Three spacers targeting exon 1, including one directly targeting the start codon, were designed using the CRISPOR tool (Concordet and Haeussler, 2018) (Fig. 5a). To generate vectors with two spacers each, single guide RNA with spacer 1 (sgRNA1) was cloned into pTRV2 vectors together with either sgRNA2 or sgRNA3 (Fig. 5b). *Cas9*-expressing explants (Skaliter et al., 2024) were inoculated with *Agrobacterium* harboring either pTRV2-*sgRNA1*-*sgRNA2* or pTRV2-*sgRNA1*-*sgRNA3* (Fig. 5b). Three independent T2 homozygous *eobv*-knockout lines were selected for further analyses: *eobv*-#1 with a 128-bp deletion between the target sites of spacers 1 and 2; *eobv*-#2 harboring two thiamine insertions, one at the spacer 1 target site and one at the spacer 2 target site; and *eobv* -#3 with a thiamine insertion at the spacer 1 (2 bp upstream of the start codon) target site and a thiamine deletion at the spacer 3 target site (Fig. 5b). In *eobv*-#1, the start codon was deleted together with the first 42 amino acids, and in lines *eobv*-#2 and #3, the mutations led to premature stop codons after 58 and 91 amino acids, respectively (Fig. 5c). Plants of the three *eobv*-knockout lines developed and flowered normally (Fig. 5d).

**Fig. 5.**
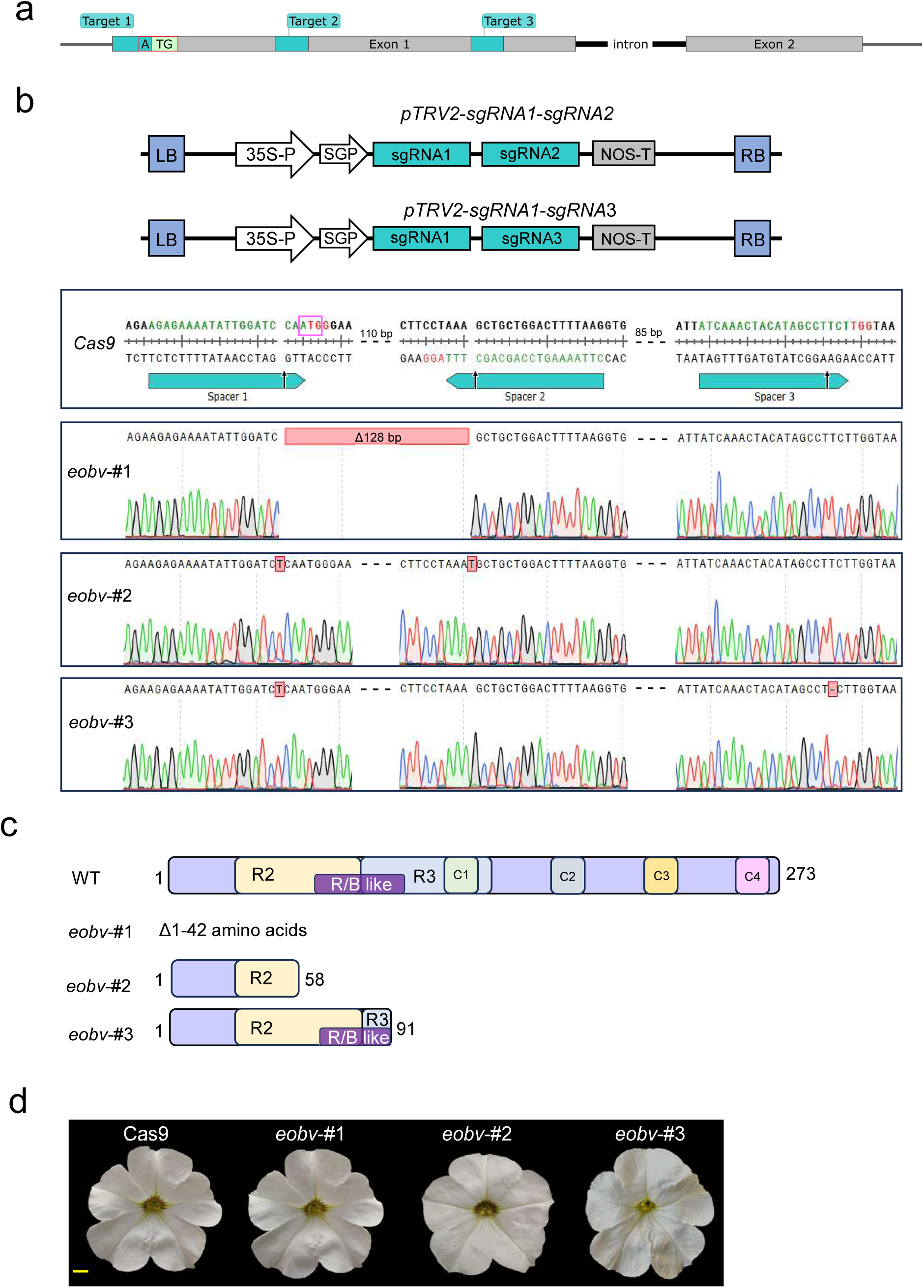
Generation of *eobv*-knockout lines using a virus-based Cas9 system. **a)** Schematic representation of the genomic sequence of *EOBV* and location of the three Cas9 target sites. **b)** Top: schematic representation of the T-DNA carrying tobacco rattle virus RNA2 (pTRV2) vectors used used for gene editing. SGP, subgenomic promoter; LB, left border; 35S-P, cauliflower mosaic virus 35S promoter; SGP, subgenomic promoter; sgRNA, single guide RNA; NOS-T, nopaline synthase terminator; RB, right border; Cas9, human co-don-optimized *Cas9* gene. Bottom: targeted genomic sequences and Sanger sequencing of *eobv*-knockout lines. Green nucleotides, spacer sequences; red nucleotides, protospacer adjacent motif (PAM). Black arrows mark the predicted cleavage site of Cas9. Magenta box, *EOBV* start codon. **c)** Predicted EOBV protein product in *eobv*-knockout and wild-type (WT) lines. Functional domains are indicated by colored rectangles. Numbers denote amino acid positions. **d)** Representative petunia cv. Mitchell flowers 2 days postanthesis from control *Cas9* and *eobv* lines. Bar = 1 cm.

To evaluate the effect of *eobv* knockout on the emission levels of volatiles, dynamic headspace analyses were conducted on flowers of *eobv*-knockout lines and control *Cas9*. In agreement with the VIGS experiment, emission levels of the C6-C3 branch compounds vanillin, eugenol and isoeugenol isomers significantly increased in *eobv*-knockout lines compared to control *Cas9*-expressing plants (Fig. 6a). In contrast, emission levels of all C6-C1/C2 compounds were significantly reduced in *eobv*-knockout lines, excluding benzyl benzoate, leading to a decrease in total emission levels in these lines (Fig. 6a). To examine whether *eobv* knockout affects the production of volatiles, internal pools were analyzed in flowers of the mutant lines and compared to control *Cas9* plants. In agreement with the headspace results, compounds generated via the phenylpropene branch (C6-C3)—isoeugenol isomers, eugenol and vanillin—significantly increased in *eobv* lines compared to control *Cas9* flowers (Fig. 6b). Levels of methyl benzoate, benzaldehyde, benzyl benzoate and benzyl alcohol (C6-C1) and phenylethyl alcohol and phenylacetaldehyde (C6-C2) were not affected by *eobv* knockout (Fig. 6b). Taken together, these results indicated that EOBV is a negative regulator of phenylpropene production and emission.

**Fig. 6.**
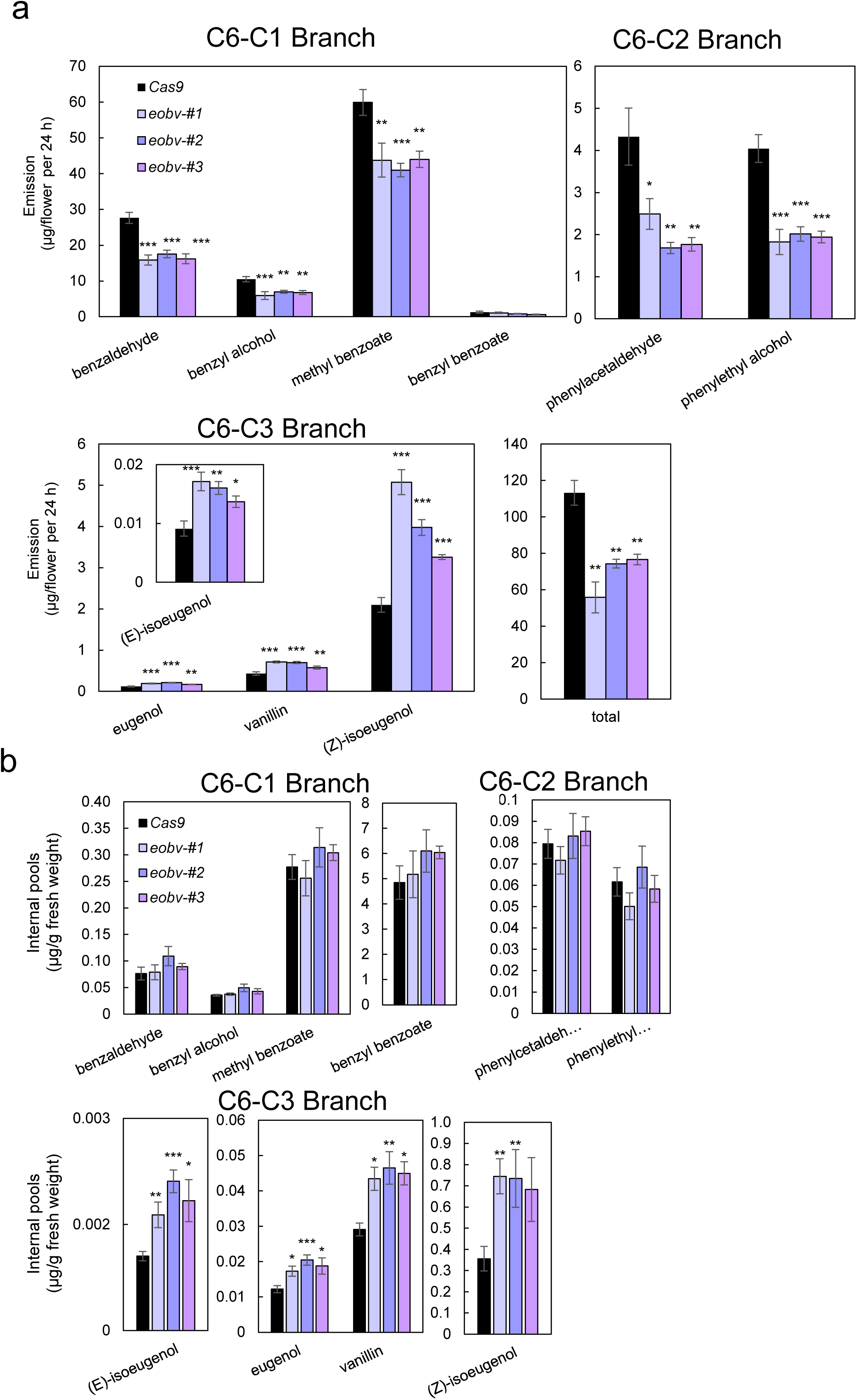
*eobv* knockout leads to an increase in phenylpropene production. **a)** Dynamic headspace analyses of *eobv*-knockout lines and *Cas9* control were performed for 24 h on flowers 2 days postanthesis (2DPA) followed by GC–MS analyses. Data are means ± SEM (n = 8–11). **b)** Internal pools were extracted from 2DPA flowers of *Cas9* (control) and *eobv* lines #1–#3 at 2000 h followed by GC–MS analysis. Data are means ± SEM (*n* = 11–15). Significance of differences was calculated using Dunnett’s or Steel’s test (\**P* ≤ 0.05, \*\**P* ≤ 0.01, \*\*\**P* ≤ 0.001) with *Cas9* as the control following one-way ANOVA. Standard errors are indicated by vertical lines.

To gain a deeper understanding of EOBV’s mode of action, transcript levels of genes encoding key biosynthesis enzymes and transcriptional regulators of volatile production were analyzed by RT-qPCR in *eobv*-knockout lines. Most of the tested genes, such as *ODO1*, *EOBI*, *EOBII*, *PhMYB4*, *EVER*, *PHENYLPROPANOID EMISSION-REGULATING SCARECROW-LIKE (PES)*, *PhDEFICIENS (PhDEF)* and *PAL1* and *2*, showed no differences in transcript levels between flowers of *eobv*-knockout lines and control *Cas9* (Fig. 7). Similarly, despite the observation that MYBs participating in the MBW complex can affect the expression of their counterparts—e.g., *AN1* expression is enhanced in leaves ectopically expressing *AN2* and in *MYB27*-RNAi lines (Spelt et al., 2000; Albert et al., 2014), transcripts levels of *AN1* and *AN11* were unaltered in *eobv* lines. Concordant with the decreased emission of phenylacetaldehyde, phenylethyl alcohol and methyl benzoate, *PHENYLACETALDEHYDE SYNTHASE* (*PAAS*) and *S-ADENOSYL-L-METHIONINE:BENZOIC ACID/SALICYLIC ACID CARBOXYL*

**Fig. 7.**
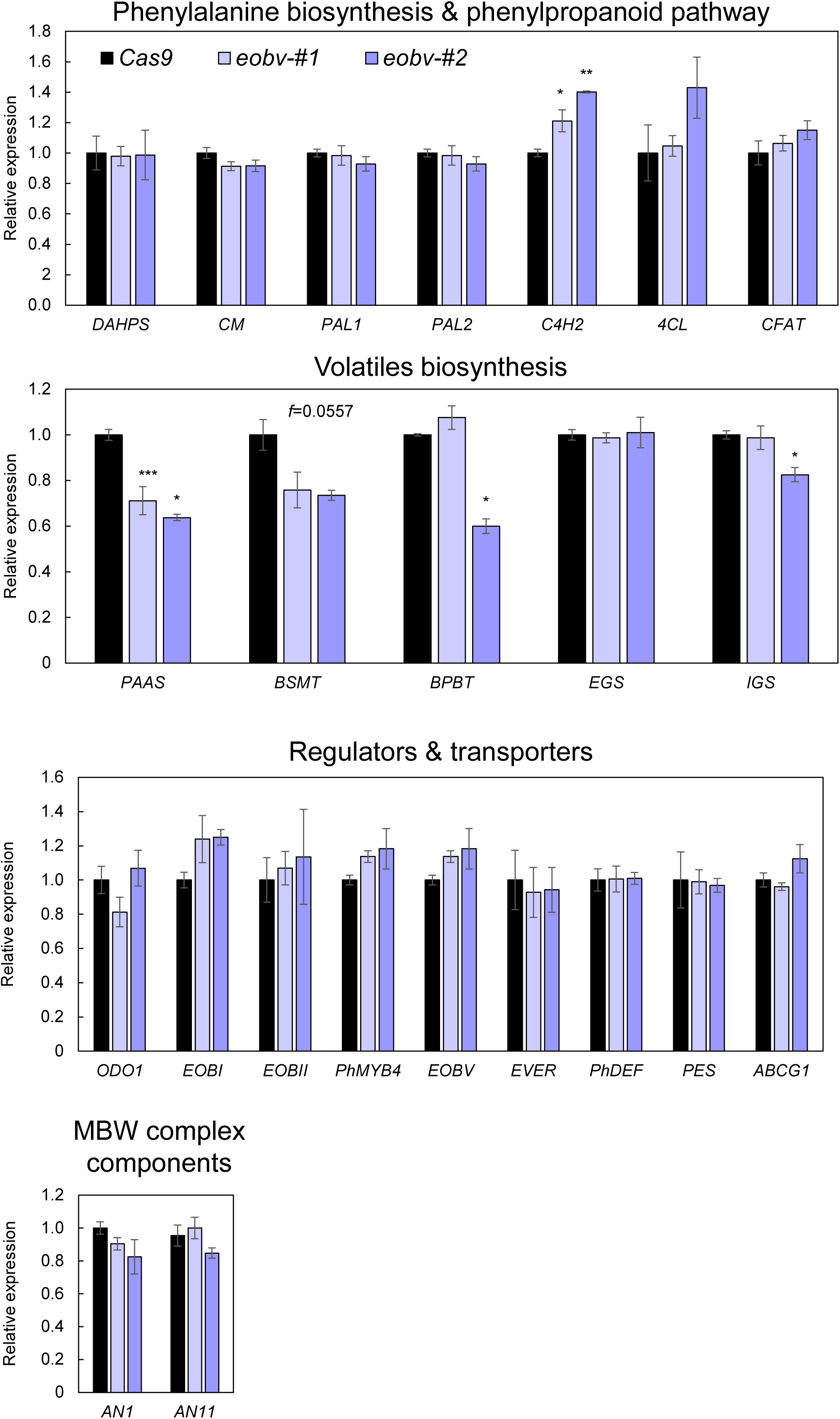
*eobv* knockout leads to an increase in *C4H* transcript level. Flowers 2 days postanthesis from *eobv*-knockout lines #1 and #2 and *Cas9* control were harvested at 1900 h and subjected to RT-qPCR analyses. Data are means ± SEM (*n* = 5). Data were normalized to the geomean of *EF1α* and *ACTIN.* Significance of differences was calculated using Dunnett’s or Steel’s test (\**P* ≤ 0.05, \*\**P* ≤ 0.01, \*\*\**P* ≤ 0.001) with *Cas9* as the control following one-way ANOVA. Standard errors are indicated by vertical lines. Phenylalanine biosynthesis and phenylpropanoid pathway genes: DAHPS, ARABINOHEPTULOSONATE 7-PHOSPHATE SYNTHASE; CM, CHORISMATE MUTASE; PAL1, PHENYLALANINE AMMONIA-LYASE 1; PAL2, PHENYLALANINE AMMONIA-LYASE 2; C4H2, CINNAMATE 4-HYDROXYLASE 2; 4CL, 4-COUMARATE:COA LIGASE; CFAT, CONIFERYL ALCOHOL ACYLTRANSFERASE. Volatile biosynthesis genes: PAAS, PHENYLACETALDEHYDE SYNTHASE; BSMT, S-ADENOSYL-L-METHIONINE:BENZOIC ACID/SALICYLIC ACID CARBOXYL METHYLTRANSFERASE; BPBT, BENZOYL-COA:BENZYL ALCOHOL/PHENYLETHANOL BENZOYLTRANSFERASE; EGS, EUGENOL SYNTHASE; IGS, ISOEUGENOL SYNTHASE. Regulator and transporter genes: ODO1, ODORANT1; EOBI, EMISSION OF BENZENOIDS I; EOBII, EMISSION OF BENZENOIDS II; EOBV, EMISSION OF BENZENOIDS V; EVER, EPIDERMIS VOLATILE EMISSION REGULATOR; PhDEF, DEFICIENS; PES, PHENYLPROPANOID EMISSION-REGULATING SCARECROW-LIKE; ABCG1, ATP-BINDING CASSETTE SUBFAMILY G MEMBER 1.

*METHYLTRANSFERASE* (*BSMT*) transcript levels decreased in the knockout lines (Fig. 6). Moreover, transcript levels of *C4H2*, which controls carbon flux to the C6-C3 branch, were significantly elevated in *eobv*-knockout lines, corresponding to the increase in emission and pools of compounds derived from that branch (Figs. 6a, 6b and 7). Transcript levels of *EOBV* were unaltered in the *eobv* lines. AtMYB4, an *Arabidopsis* homolog of EOBV, has been shown to repress its own promoter, hence knockout of *eobv* may lead to a break in a negative feedback loop (Zhao et al., 2007). Furthermore, the system’s ability to recognize aberrant mRNAs is highly complex and requires multiple factors to trigger RNA degradation via nonsense-mediated mRNA decay (Luha et al., 2024).

The promoter of *C4H2* contains two repeats of the MYB binding motif YACCWACY at nucleotide positions –115 and –205. AtMYB4 has been shown to bind to this motif in promoter regions proximal to the transcription start site (Agarwal et al., 2020). In contrast to *C4H2*, *PAAS* and *BSMT* promoters do not contain this motif (Fig. S9a). To identify additional potential targets of EOBV, in-house Python-based motif scanning of the petunia genome was performed, as bioinformatic tools such as MEME (meme-suite.org) do not include a petunia database. To this end, 300-bp sequences from all annotated genes in the *P*. *axillaris* genome (Bombarely et al., 2016) were extracted and scanned for the motif YACCWACY (e value ≤ 0.005). Genes with low or no expression in the petunia petal transcriptome (Shor et al., 2023b) or lacking annotation were excluded. Moreover, as multiple motifs are often required for efficient regulation of target genes, as also evident from the two copies present in the *C4H2* promoter, only sequences containing two or more copies of the motif were included (Espley et al., 2009); 39 candidate genes were identified. According to predicted annotations, these candidates are involved in lignin and phenylpropanoid biosynthesis, cell wall remodeling and expansion, hormonal signaling, and secondary metabolism, integrating structural, developmental, flowering and stress-related processes, and representing a broad range of processes potentially involving EOBV (Table S1). Among the genes involved in phenylpropanoid biosynthesis was another copy of *C4H* in petunia—*C4H1*, whose expression is much lower in petunia flowers when the scent machinery is active compared to that of *C4H2* (Colquhoun et al., 2011). In addition, two genes involved in aromatic amino acid biosynthesis were identified: *CHORISMATE SYNTHASE (CS)* responsible for conversion of 5-enolpyruvylshikimate-3-phosphate to chorismate and *AROGENATE DEHYDRATASE 3* (*ADT3*), which converts arogenate to phenylalanine, the main precursor of phenylpropanoids. To determine whether EOBV regulates these genes, their expression was analyzed in *eobv* flowers by RT-qPCR. Transcript levels of *ADT3* were significantly higher in *eobv* compared to control *Cas9* flowers (Fig. S9b). In contrast, transcripts of *C4H1* and *CS* were unaltered in the *eobv* flowers. These findings indicated that EOBV regulates genes encoding enzymes involved in both the synthesis of phenylpropanoid precursors and the downstream steps of the phenylpropanoid pathway.

The phenotypes observed in the *eobv*-knockout lines, i.e., increased emission of phenylpropenes with concomitant decrease in benzenoids and elevated levels of *C4H2*, resemble the effect reported by Colquhoun et al. (2011) for RNAi-mediated suppression of *PhMYB4*. However, although the diel pattern was similar, the spatial developmental pattern was different: in contrast to EOBV, *PhMYB4* was not enriched in the adaxial epidermis (Fig. S10). Moreover, in the *eobv*-knockout lines, *PhMYB4* transcript levels were similar to those in the *Cas9* control, indicating that EOBV does not affect *PhMYB4* expression (Fig. 7). Although off-target gene-editing events induced by Cas9 in plants are generally considered to be rare, they have been reported to occur at genomic sites with high sequence similarity to the target (Sturme et al., 2022). Considering a nucleotide identity between *EOBV* and *PhMYB4* of 80.34% (mRNA), the three spacers selected to target *EOBV* contained 4 to 10 mismatches relative to *PhMYB4*, including 2 to 5 mismatches within the critical seed sequence, to lower the chances of off-target gene editing (Fig. S11a). Sanger sequencing of *PhMYB4* in *eobv*-knockout lines confirmed that no editing events had occurred in *PhMYB4*’s sequences in those lines (Fig. S11b). Taken together, these results indicated that the phenotype of *eobv*-knockout lines is not dependent on PhMYB4.

### Stacking of *ever-* and *eobv*-knockout alleles has no additive effect on phenylpropene emission in petunia flowers

Given their major relevance in the ornamental, agricultural, food and beverage, and pharmaceutical industries, enhanced production of phenylpropene volatile compounds is a biotechnological goal (Skaliter et al., 2022). We previously identified EVER, a MYB transactivator that regulates the emission of low-vapor-pressure volatiles, e.g., phenylpropenes, by modulating the composition of petal epicuticular waxes (Skaliter et al., 2024). Because EVER and EOBV influence phenylpropene levels through distinct mechanisms—modification of the epicuticular wax layer (Fig. 1) and suppression of *C4H2* and *ADT3*, respectively—we tested whether generating petunia flowers with double knockouts of *ever* and *eobv* would result in an additive increase in phenylpropene emission; *ever* line #2, which exhibited an increase of ca. 1.7-fold in emission levels of phenylpropenes (Skaliter et al., 2024), and *eobv* line #1 were cross-pollinated to produce F1 hybrids that were subsequently self-pollinated until an F3 population with homozygous knockout mutations was generated. F3 plants and flowers of *ever* x *eobv* developed normally and did not differ from *eobv*, *ever* or control *Cas9* (Fig. S12a). Headspace analysis revealed that flowers of *ever* x *eobv* had higher levels of phenylpropenes than the control *Cas9*; however, none of the volatiles were at higher levels than in the parental lines *eobv* and *ever* (Fig. S12b), suggesting limited carbon availability for phenylpropanoid volatile production.

### Knockout of *eobv* mitigates the effect of high ambient temperature on floral traits

*EOBV* transcripts are heat-responsive, their levels significantly increasing in flowers of petunia cvs. P720 and Blue Spark grown at high ambient temperature (Cna’ani et al., 2014). To detail the involvement of EOBV in regulating volatiles under high ambient temperature, plants of *eobv*-knockout lines and control *Cas9* were first grown under standard growth conditions (22/16°C day/night) and then half were transferred to a higher temperature regime (28/22°C day/night). In line with Sood et al. (2021), the control flowers were significantly affected (∼12% smaller in diameter) by the high ambient temperature (Fig. 8a). As expected, there was no difference in diameter between flowers of *Cas9* and *eobv* at 22/16°C (Figs. 5d and 8a). High temperature had a less pronounced effect on flower size of the *eobv* plants (6% reduction) and their diameter remained higher than those of *Cas9* plants grown at 28/22°C (Fig. 8a). RT-qPCR analysis targeting *EOBV* transcripts revealed an increased level in flowers of *Cas9* plants grown at higher ambient temperature, confirming that *EOBV* is also heat-responsive in the ‘Mitchell’ genomic background (Fig. 8b). *EOBV* transcript levels were unaltered in *eobv* knockouts at elevated temperature, indicating that the multicomplex machinery regulating accumulation of transcripts does not recognize aberrant *eobv* mRNA under both standard and heat-stress conditions. To assess whether the difference in petal size observed under elevated temperatures is associated with changes in α-expansin (PhEXP1), shown to be involved in regulation of petal size (Zenoni et al., 2004), its transcript levels were quantified in *Cas9* and *eobv* flowers grown at 22/16°C and 28/22°C. *PhEXP1* expression was significantly reduced at 28°C in both genotypes to a similar extent, indicating that its downregulation under heat conditions occurs independently of EOBV (Fig. S13).

**Fig. 8.**
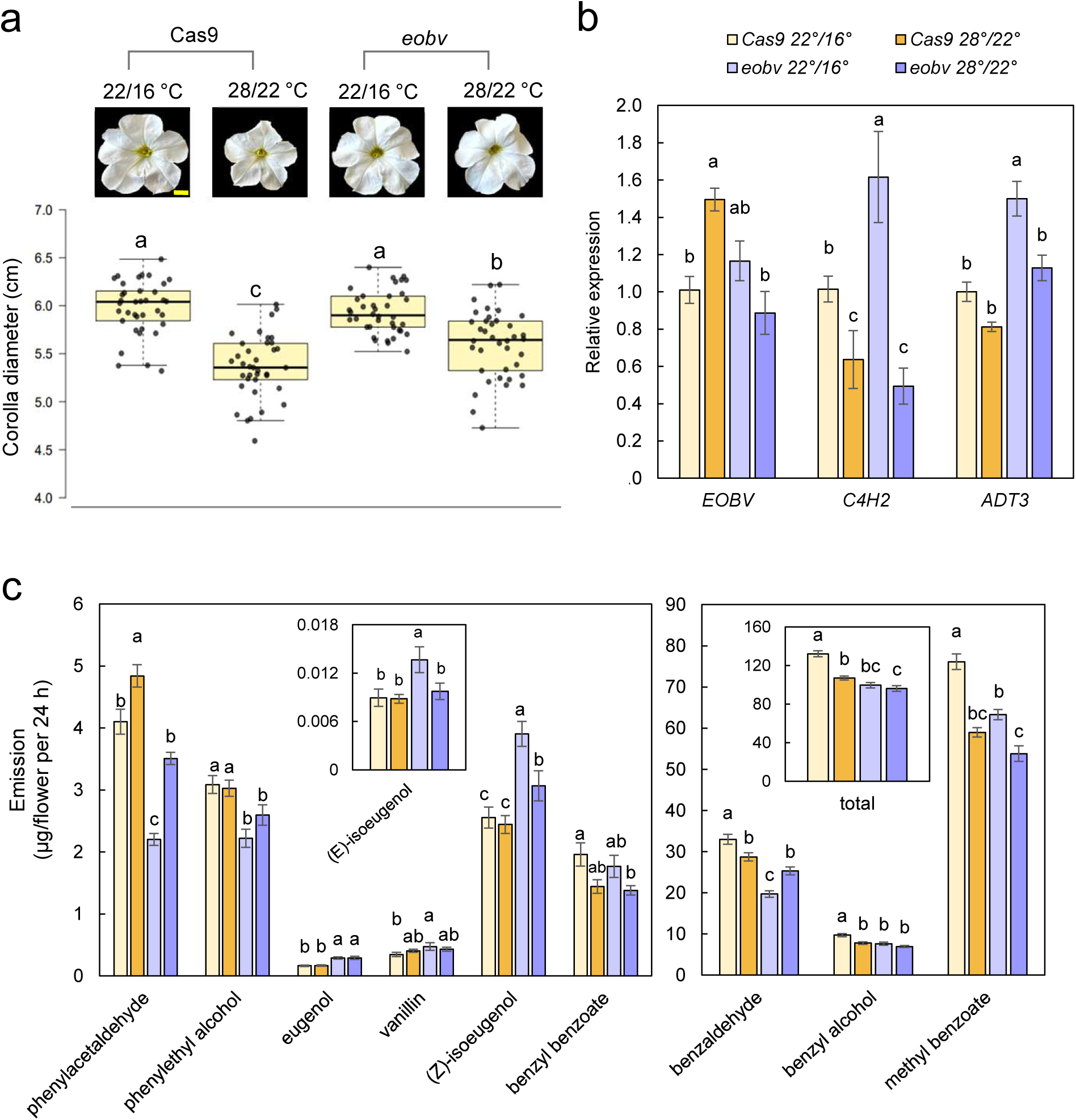
Effect of elevated growth temperature regime on *eobv*-knockout lines. Plants were grown for 1 month in a phytotron at 22/16°C or 28/22°C. **a)** Top: representative images of 2 day postanthesis (2DPA) flowers of the different lines and regimes. Bar = 1 cm. Bottom: Diameter of corollas from the different treatments (*n* = 38). **b)** RT-qPCR analysis of RNA extracted from flowers 2DPA. Data are means ± SEM (*n* = 5). Data were normalized to the geomean of *EF1α* and *ACTIN.* **c)** Dynamic headspace analysis was performed on flowers 2DPA for 24 h, followed by GC–MS analysis. Data are means ± SEM (*n* = 10–16). Significance of differences was calculated by Tukey’s multiple comparison test following one-way ANOVA. Values with different letters are significantly different at *P* ≤ 0.05. Standard errors are indicated by vertical lines. C4H2, CINNAMATE 4-HYDROXYLASE 2; ADT3, AROGENATE DEHYDRATASE 3.

Analysis of expression levels of EOBV’s target, *C4H2*, revealed that it is significantly reduced in response to high temperature in both *Cas9* and *eobv* flowers (Fig. 8b). Moreover, transcript levels of *PhMYB4*, which also regulates *C4H2*, decreased slightly, albeit not significantly, at higher ambient temperatures in both *eobv* and *Cas9* (Fig. S13). Expression levels of *ADT3*, the other target of EOBV, were only mildly affected by heat stress, showing a non-significant decline in *Cas9* flowers grown at 28/22°C compared to 22/16°C. In *eobv* lines, *ADT3* transcript levels decreased significantly under elevated temperature, reaching levels comparable to those in *Cas9* flowers. Dynamic headspace analysis of *eobv* and *Cas9* flowers revealed that, consistent with Cna’ani et al. (2015), total volatiles in *Cas9* at 28/22°C are significantly reduced, owing to the decrease in the major emitted compounds—benzaldehyde, benzyl alcohol and methyl benzoate (Fig. 8c). In *eobv* flowers, total emission levels were not significantly altered, because levels of methyl benzoate decreased, benzyl alcohol levels remained unchanged, and benzaldehyde levels increased. Interestingly, the emission levels of phenylacetaldehyde were significantly increased at 28/22°C in both *Cas9* and *eobv* lines. None of the phenylpropenes’ levels were affected by high ambient temperature in *Cas9* flowers, whereas isoeugenol isomers in *eobv* declined significantly, revealing that carbon flux under high temperature is affected by EOBV but probably not via C4H2 or ADT3.

## Discussion

Specialized metabolite biosynthesis is not confined to individual pathway branches but occurs through interconnected activity across multiple metabolic layers. For example, increasing flux through the phenylpropanoid pathway led to enhanced terpenoid biosynthesis, and vice versa (Zvi et al., 2012; Skaliter et al., 2026). At the inter-pathway level of phenylpropanoid metabolism, anthocyanin and volatile biosynthesis are tightly interlinked, as shifting metabolic flux toward either branch can enhance production in the other, and in some cases, increased flux leads to simultaneous enhancement of both branches (Zuker et al., 2002; Zvi et al., 2008; Zvi et al., 2012; Cna’ani et al., 2014; Cna’ani et al., 2015; Shaipulah et al., 2016; Ravid et al., 2017). Conversely, high ambient temperatures lead to a decrease in both anthocyanins and volatile production (Cna’ani et al., 2014; Sood et al., 2021). Indeed, several transcription factors—MYB PH4, PRODUCTION OF ANTHOCYANIN PIGMENT 1 and PhDEF—have been shown to control both pigments and scent (Quattrocchio et al., 2006; Zvi et al., 2008; Zvi et al., 2012; Cna’ani et al., 2015). In this study, we reveal an additional element that regulates these showy traits—the petunia MBW complex. VIGS suppression of the MBW components *AN1* (bHLH) or *AN11* (WDR) in petunia flowers resulted in increased emission of several volatiles from the three branches of phenylalanine-derived volatiles, including a 2-fold rise in methyl benzoate, the main volatile emitted by ‘Mitchell’ flowers. These results suggest that the MBW complex has a broad negative regulatory role in volatile production. The activity and targets of MBW complexes are dictated by the MYB component (Albert et al., 2011; Albert et al., 2014). Therefore, interactions between volatile-related MYB factors and AN1 and AN11 were tested. Y2H and in-planta pairwise assays revealed that the volatile MYB regulator EOBV interacts directly with AN1. Notably, EOBV, like its *Arabidopsis* homolog AtMYB4 which regulates anthocyanins and interacts with bHLH members, also has the conserved R/B-like bHLH binding motif (Wang et al., 2020). Three-way in-planta protein–protein interaction assays confirmed the formation of the MBW complex consisting of EOBV, AN1 and AN11. MBW components from different plant species have been shown to complement one another (Quattrocchio et al., 1993; de Vetten et al., 1997; Albert et al., 2011). Indeed, ectopic expression of only LacI-AN1 in the pairwise assay, and of LacI-EOBV/AN11-GAL4 and LacI-EOBV/AN1 in the three-way assays, led to weaker activation of the reporter gene in *N*. *benthamiana* leaves, likely due to the presence of heterologous MBW components.

Spitzer-Rimon et al. (2012) suggested a negative effect of EOBV on floral scent production, although its exact function and mode of action remained unknown. Detailed analyses of volatiles in *eobv*-knockout flowers suggest that EOBV fine-tunes phenylpropene levels: these were significantly increased in the mutant lines relative to control *Cas9* flowers. In contrast, the pools of benzenoids and phenylpropanoid-related compounds was unaltered while their emission decreased, emphasizing the complexity of the machinery responsible for the biosynthesis, intracellular accumulation, transport and release of volatiles (Adebesin et al., 2017; Cna’ani et al., 2017; Liao et al., 2021; Liao et al., 2023; Skaliter et al., 2024). Similar to the *eobv*-suppressed/*eobv*-knockout lines, *AN1*/*AN11* suppression resulted in increased phenylpropene emission. Yet, the emission of C6-C1/C2 volatiles increased in *AN1*/*AN11*-suppressed tissues while their levels were reduced in *eobv*-suppressed tissue. These results indicate that AN1 and AN11 exert a broader influence on volatile production, through direct or indirect mechanisms that are independent of EOBV, potentially by engaging other MYB partners. For example, AN1 and AN11 are necessary for the production of flavonoids, which have been shown to negatively regulate the expression of key phenylpropanoid pathway genes such as *PAL* (Yin et al., 2012). Therefore, suppression of *AN1* or *AN11*, or both, may shift the metabolic balance and alleviate this negative feedback, potentially restoring or enhancing upstream gene expression. Moreover, AN1 has been shown to be involved in the perception/transduction of ethylene signal (Prinsi et al., 2016), and ethylene has been shown to negatively regulate volatiles in petunia (Liu et al., 2017). Notably, the difference in volatile profiles observed following suppression of AN1 or AN11, i.e., the increase in (Z)-isoeugenol detected only in the latter, may be ascribed to the function of WDR proteins as scaffolds anchoring different bHLHs: in petunia, AN11 interacts with JAF13, in addition to AN1 (Albert et al., 2014); in *Arabidopsis*, the WDR TRANSPARENT TESTA GLABRA1 is essential for trichome formation, seed coat pigmentation, and anthocyanin production, with no single bHLH being able to compensate for its loss; in contrast, individual bHLHs such as GLABRA3 or ENHANCER OF GLABRA3 are partially redundant, and their loss typically produces only mild or tissue-specific phenotypes (Ramsay and Glover, 2005).

Transcript levels of *C4H2*, encoding the enzyme that controls carbon flux toward phenylpropene biosynthesis, were enhanced in flowers of *eobv* knockouts, in line with phenylpropenes’ emission and pool results. Based on our in silico promoter analysis, *ADT3* was also upregulated in *eobv* knockouts, but probably does not have a major impact on total carbon flux as this is not the main *ADT* expressed in mature petunia flowers (Maeda et al., 2010). The promoters of *C4H2* and *ADT3* each contain two repeats of a more specific version of the YACCWACY consensus sequence, which starts with a cytosine and is therefore defined as CACCWACY. Interestingly, several genes carrying only a single copy of CACCWACY (with the exception of *PAL2*): *CS*, *CM*, *PAL1* and *C4H1*, did not show altered expression in the *eobv*-knockout lines. Further experimental analysis will determine whether two CACCWACY elements are required for efficient EOBV-mediated repression of target genes. In contrast to *C4H2* and *ADT3*, which were upregulated in *eobv*-knockout lines, *BSMT* and *PAAS*, which were downregulated, do not contain the YACCWACY motif in their promoters. Because EOBV is a repressor, their downregulation in *eobv* is more likely due to loss of repression of their transcriptional repressors or repression of their activators, or alternatively, due to a feedback loop that prevents toxicity due to hyperaccumulation of volatile organic compounds (Liao et al., 2021; Liao et al., 2023).

Despite the facts that *EOBV* and *PhMYB4* have similar diel expression patterns, both regulate *C4H2* and their suppression has a similar effect on volatile emissions, several lines of evidence differentiate them: (1) *ADT3* transcripts were upregulated only in *eobv*-knockout lines; (2) their developmental expression pattern is different (Figs. 5 and 8a); (3) *EOBV* transcripts are enriched in the adaxial epidermis, whereas those of *PhMYB4* are not (Figs. 4d and S10b); (4) knockout of *eobv* had no effect on transcript levels of *PhMYB4*; (5) *PAL* and *C4H1* transcripts were higher in *PhMYB4-*RNAi lines, whereas in the *eobv* knockout, *PAAS* and *BSMT* expression was lower (Colquhoun et al., 2011); (6) EOBV interacted with AN1, but PhMYB4 did not; (7) *PhMYB4* transcripts did not respond to high ambient temperature, in contrast to *EOBV*. Together with analysis of the genomic context (Bombarely et al., 2016), it can be suggested that *EOBV* and *PhMYB4* are paralogs that have probably undergone subfunctionalization. Further experiments are needed to corroborate the exact relations between these two regulators.

In addition to spatial, developmental and diel regulation, *EOBV* expression responds to high ambient temperatures. Characterization of *eobv*-knockout lines under long-term exposure to elevated growth temperatures revealed that EOBV is also involved, albeit to a lower extent, in regulating scent production under heat-stress conditions. Transcript levels of EOBV’s target, *C4H2*, declined to a similar extent in both *eobv* and *Cas9* flowers under elevated temperatures, regardless of *EOBV*‘s increased expression. Furthermore, the observation that *PhMYB4*, which also regulates *C4H2*, remains unaffected by heat suggests that it is also not involved in heat regulation of *C4H2*. Levels of the other EOBV target, ADT3, were slightly, but not significantly reduced in heat-stressed flowers of *Cas9* plants, indicating that it is only mildly affected by elevated temperatures. A similar trend was observed for *ADT3* in *eobv* flowers; however, the reduction was significant, and transcript levels reached those of *Cas9* plants. This likely reflects the higher basal expression of *ADT3* in the knockout lines, rendering it more susceptible to heat-induced downregulation. Cna’ani et al. (2015) showed that in petunia plants grown under elevated temperature, transcript levels of genes encoding enzymes for aromatic amino acid biosynthesis, as well as enzymes dedicated to volatile production, including *C4H*, are drastically reduced, indicating a global plant response aimed at limiting phenylpropanoid biosynthesis. Furthermore, extensive transcriptomic reprogramming that activates the heat-stress response was shown in plants exposed to high temperatures (Chen and Li, 2017). Taken together, these results suggest that EOBV does not mediate the heat regulation of *C4H2* or *ADT3*, and that both are regulated as part of a broader temperature-responsive network.

Heat is a key environmental factor that has been shown to negatively impact flower size in various plants (Descamps et al., 2018; Sood et al., 2021; Wiszniewski et al., 2022). In this study, we show that EOBV is involved in determining final flower size under heat-stress conditions; the reduction in corolla size caused by high temperatures was mitigated in *eobv*-knockout compared to control *Cas9* plants. Although *EOBV* transcripts are enriched in the adaxial epidermis under standard growth conditions, there was no differential effect on the development of adaxial versus abaxial sides, as overall floral morphology remained normal at high temperature. It would be informative to test *EOBV* bilateral expression in the petal epidermis at different stages of flower development, especially under high-temperature conditions, as well as its post-transcriptional/translational regulation. It is worth noting that *cycloidea* mutants in *Antirrhinum majus* (Snapdragon) have smaller petals although, like EOBV, *CYCLOIDEA* is expressed predominantly in the adaxial epidermis (Luo et al., 1996).

Flower cells undergo proliferation and expansion during the early and late stages of bud development, respectively, both of which have been shown to be impacted by elevated temperatures (Sood et al., 2021). Indeed, expression of petunia *PhEXP1*, which plays a key role in cell expansion, was reduced in flowers exposed to high temperature (Fig.13) (Zenoni et al., 2004). However, its expression was similarly reduced in both *eobv* and *Cas9* flowers at 28°C, suggesting that the limited reduction in petal size observed in *eobv* under elevated temperatures was not due to misregulation of *PhEXP1*, and that the effect on flower size may be mediated by additional factors downstream or independent of PhEXP1. Sood et al. (2021) demonstrated that in petunia, increased levels of abscisic acid affect early stages of bud development in response to elevated temperatures and likely antagonize cytokinin-promoted cell division. Interestingly, crosstalk between abscisic acid and the MBW complex has been demonstrated in several plants (Lama et al., 2020; Broucke et al., 2023). Moreover, our in silico promoter analysis revealed that the promoters of several genes linked to hormones contain repeats of the YACCWACY motif, found in targets of EOBV. Two repeats of this motif were also found in a homolog of the *Arabidopsis* MADS-box transcription factor *SHORT VEGETATIVE PHASE*, a heat-sensitive repressor of *FLOWERING LOCUS T*; mutations in the latter have been shown to result in smaller petals (Jeong et al., 2007; Romera-Branchat et al., 2025). Thus, at early stages, EOBV might modulate the expression of genes related to flower initiation/development, and this modulation could become dysregulated under heat stress, leading to changes in petal size. The interaction of EOBV with the components of the MBW complex, involved in the control of floral showy traits, further interlinks flower development with the environment.

## Materials and methods

### Plant material and growth conditions

*Petunia × hybrida* cv. Mitchell diploid plants were grown from seeds. Rooted plants of *Petunia × hybrida* cv. Classic Blue Ray were obtained from Danziger – “Dan” Flower Farm (Mishmar Hashiva, Israel). Plants were grown in a greenhouse under 25°C day/20°C night temperatures with a 16 h light/8 h dark photoperiod [incandescent light bulbs (120 V, 16 lm/W) were used to provide additional lighting].

For the elevated growth temperature experiment, *eobv*-knockout line #1 and control *Cas9* seedlings were sown and grown for a month under standard conditions (25/20°C day/night). Then, plants from each treatment, which had not yet developed floral buds, were separated into two groups and transferred to a phytotron greenhouse (16/8 h light/dark photoperiod) with controlled rooms: one group from each treatment at 22/16°C day/night and the other at 28/22°C. After a month, 2DPA flowers were taken for diameter measurement with a Vernier caliper (Mitutoyo, Kanagawa, Japan), and headspace, internal pool and RT-qPCR analyses.

*N. benthamiana* plants were kindly provided by Dr. Guy Polturak (Institute of Plant Sciences and Genetics in Agriculture, The Robert H. Smith Faculty of Agriculture, Food and Environment, The Hebrew University of Jerusalem) and maintained in a controlled growth room at 22°C under a 16 h light/8 h dark photoperiod with fluorescent lighting.

### RNA extraction and RT-qPCR analyses

Total RNA was extracted from 30–100 mg of ground (with liquid nitrogen) floral tissues using the Tri-Reagent kit (Sigma-Aldrich) and treated with RNase-free DNase I (Thermo Fisher Scientific). First-strand cDNA was synthesized using total RNA, oligo(dT) primer, and reverse transcriptase ImProm-II (Promega, Madison, WI, USA) according to the manufacturer’s instructions. Two-step real-time qPCR was performed on a CFX Opus 384 Real-Time PCR System (Bio-Rad, Hercules, CA, USA) using 2X qPCRBIO SyGreen Blue Mix Hi-ROX (PCR Biosystems, London, UK). A standard curve was generated for each gene using dilutions of cDNA samples, and data analysis was performed using Bio-Rad CFX Maestro software (Bio-Rad, Hercules, CA, USA). Primer specificity was determined by melting-curve analysis. Raw transcript-level data were normalized to the geomean of *EF1α*, *UBIQUITIN* and *ACTIN*. Quantification calculations were carried out using the 2^−ΔΔCT^ formula. Primers are shown in Table S2.

### VIGS suppression of target genes in petunia flowers

A 162-bp fragment of the 3’ UTR of *EOBV*, a 332-bp fragment of AN11’s 3’ UTR and a 250-bp fragment of AN1’s 3’ UTR were amplified and cloned into pTRV2. *Agrobacterium tumefaciens* strain AGL0 was transformed with plasmids harboring RNA1 (pTRV1), pTRV2-EOBV, pTRV2-AN1, pTRV2-AN11 or pTRV2-CHS as a control. *Agrobacterium* carrying pTRV1 were mixed at a 1:1 ratio (to OD_600_ = 0.5) with *Agrobacterium* carrying the above-mentioned plasmids in inoculation solution containing 200 μm acetosyringone and 10 mm MgCl_2_. Corollas of petunia cv. Mitchell flowers at anthesis were inoculated with the *Agrobacterium* solutions by piercing with a needle and infiltration using a syringe, followed by localized headspace analysis as described in Skaliter et al. (2021).

To determine EOBV’s effect on anthocyanin content in petunia cv. P720, cultures of *Agrobacterium* transformed with pTRV1 and pTRV2-EOBV or pTRV2-CHS were grown overnight at 28°C in LB Broth Lennox medium (Formedium, Norfolk, UK) supplemented with 50 mg/L kanamycin and 200 µM acetosyringone. The cells were harvested and resuspended in inoculation buffer containing 10 mM MES, pH 5.5, 200 µM acetosyringone, and 10 mM MgCl_2_ to an OD_600_ of 10. Following an additional 3 h of incubation at 28°C, the bacteria containing pTRV1 were mixed with those containing the pTRV2 derivatives in a 1:1 ratio; 400 μL of this mixture was applied to the cut surface, after removing the apical meristem, of 1-month-old petunia ‘P720’ plants.

### Collection of emitted volatiles, internal pools and GC–MS analysis

Dynamic headspace analysis was performed by harvesting flowers 2DPA and placing them in a 50-mL beaker filled with tap water in jars (2 flowers per jar). Volatiles were collected for 24 h using columns made of glass tubes containing 100 mg Porapak Q polymer and 100 mg 20/40-mesh, held in place with plugs of silanized glass wool. Trapped volatiles were eluted with 1.5 mL hexane and 0.5 mL acetone. Isobutylbenzene was used as an internal standard.

To determine the pool sizes of volatile compounds, 100 mg tissue from 1DPA flowers was ground in liquid nitrogen and extracted in 400 μL hexane containing isobutylbenzene as the internal standard. Following 2 h of incubation with gentle shaking at 25 °C, extracts were centrifuged at 10,500× g for 10 min, and the supernatant was further centrifuged, followed by GC–MS analysis.

GC–MS analysis (of a 1 μL sample) was performed using a device consisting of a PAL autosampler (CTC Analytics, Zwingen, Switzerland), a TRACE GC 2000 gas chromatograph (Thermo Fisher Scientific, Waltham, MA, USA) equipped with an Rtx-5SIL mass spectrometer fused-silica capillary column (inner diameter 0.25 μm, 30 m × 0.25 mm; Restek, Centre County, PA, USA), and a TRACE DSQ quadruple mass spectrometer (Thermo Fisher Scientific, Waltham, MA, USA). Helium was used as the carrier gas at a flow rate of 1 mL/min. The injector temperature was set to 220 °C (splitless mode), the interface to 240 °C, and the ion source to 200 °C. The analysis was performed under the following temperature program: 2 min of isothermal heating at 40 °C followed by a 7 °C/min oven temperature ramp to 250 °C, then 2 min of isothermal heating. The system was equilibrated for 1 min at 70 °C before injecting the next sample. Mass spectra were recorded at 3.15 scans/s with a scanning range of 40–350 mass-to-charge ratio and electron energy of 70 eV. Compounds were tentatively identified (>95% match) based on NIST/EPA/NIH Mass Spectral Library data version NIST 17 (with software version 3.4) using xcalibur 1.3 (Thermo Fisher Scientific, Waltham, MA, USA). Further identification of all compounds was based on a comparison of mass spectra and retention times with those of authentic standards (Sigma-Aldrich, St. Louis, MO, USA) analyzed under similar conditions.

### Protein–protein interaction assays

#### Y2H assay

The coding sequences of the tested genes were cloned to either the GAL4 BD in the pBD-GAL4 CAM vector (Clontech) or the GAL4 AD in pACT2 (Takara Bio, Mountain View, CA,USA) and transformed into *Saccharomyces cerevisiae* strain Y190.

Transformed yeast were grown on synthetic defined medium (SD) lacking leucine and/or tryptophan for selection. Interaction between targets was detected by dripping 2.5-μL drops of individual clones on SD media lacking leucine and tryptophan followed by incubation at 28°C for 48 h. Formed colonies were gently covered with chloroform and incubated for 15 min at room temperature, and then the chloroform was drained. Colonies were gently covered with staining solution (containing X-Gal) and monitored for β-galactosidase reporter gene expression levels after 1 h.

#### In-planta protein-interaction assays

All vectors were constructed using GoldenBraid 2.0 (Sarrion-Perdigones et al., 2013). The *RUBY* open-reading frame (He et al., 2020) was cloned under the 6×LacI operon fused to the CaMV 35S minimal promoter (pDGB3α1-pOpLacI:mini35S:*RUBY*). *AN1* was fused with the LacI BD into pDGB3α1 or cloned into pDGB3 3Ω1 containing *DsRed*, and *AN11* was fused with the GAL4 AD. EOBV was fused with either GAL4 AD or LacI BD into pDGB3α2. *Agrobacterium* were mixed (to OD_600_ = 0.5) and third or fourth leaves of *N*. *benthamiana* plants or *N*. *tabacum* disc explants were inoculated and placed on regeneration media. In the three-way protein–protein interaction assays, inoculated *N*. *benthamiana* leaves were examined under a fluorescent stereomicroscope (SMZ1270; Nikon, Minato City, Tokyo, Japan) for DsRed detection. Inoculated leaves of both pairwise and three-way assays were harvested 5 days postinoculation, and 3 disks of each leaf (0.8 cm diameter) were taken for analysis and immersed in 100% ethanol until the tissue was completely bleached. Betalains were extracted from the tissue by the addition of 1 mL DDW and placed on an orbital shaker (150 rpm) at room temperature for 3 h. The light absorbance of the extract was measured in a Multiskan SkyHigh Microplate Spectrophotometer (Thermo Fisher Scientific, Waltham, MA, USA) at 538 nm. For *N. tabacum*, 2 weeks postinoculation, calli formed from inoculated explants were bleached in 100% ethanol for 2 days and observed under a stereomicroscope (SMZ1270; Nikon, Minato City, Tokyo, Japan).

#### Phylogenetic analysis

Protein sequences were aligned using the Clustal W algorithm (Larkin et al., 2007) implemented in MEGA11 (Tamura et al., 2021) with default parameters (gap opening penalty 10.00; gap extension penalty 0.20). Maximum likelihood with default parameters (1000 bootstraps; Jones–Taylor–Thornton substitution model) was used as the statistical method for phylogenetic analysis in MEGA11.

#### EOBV localization assay

The coding sequence of *EOBV* (without the stop codon) was amplified from cDNA extracted from flowers and fused to GFP using a GoldenBraid 2.0 reaction (Sarrion-Perdigones et al., 2013) to generate pDGB3α2-EOBV-GFP and pDGB3α2-GFP-EOBV. ‘Mitchell’ flowers at anthesis were inoculated with a mixture of *Agrobacterium* carrying pDGB3α2-EOBV-GFP and pRCS-RFP-VirD2-NLS (marker for nuclear localization) or pDGB3α2-GFP-EOBV and pRCS-RFP-VirD2-NLS, or pDGB3α2-EOBV-GFP alone and pRCS-RFP-VirD2-NLS alone as a control. Inoculated flowers were harvested 2DPA. Images were acquired with a Leica SP8 laser scanning microscope equipped with solid-state lasers with 488 nm (GFP excitation) and 552 nm (RFP excitation) light, an HC PL APO CS 63x/1.2 water-immersion objective and Leica Application Suite X software (LASX) (Leica Microsystems, Wetzlar, Germany). GFP- and RFP-emission signals were detected with HyD (hybrid) detectors in the ranges of 500–540 and 565–640 nm, respectively. Merged images were generated using Fiji (Image J) software (Rueden et al., 2017).

#### Anthocyanin content

To determine anthocyanin levels, 100 mg of corollas of petunia cv. Classic Blue Ray inoculated with TRV2-EOBV or TRV2-CHS were ground (with liquid nitrogen), incubated with acidic methanol (containing 1% v/v HCl) and placed on an orbital shaker (150 rpm) at 4°C for 2 h in the dark. The light absorbance of the extracts was measured in a spectrophotometer (Multiskan SkyHigh spectrophotometer; Thermo Fisher Scientific, Waltham, MA, USA) at 530 nm (anthocyanins) and 657 nm (chlorophyll). Anthocyanin content was calculated using the using the formula A_530_ – 0.25(A_657_).

#### Generation of *eobv*-knockout lines

Tobacco rattle virus RNA2 (pTRV2) vectors pTRV2-sgRNA1-sgRNA2 and pTRV2-sgRNA1-sgRNA3, were synthesized by Twist Bioscience (San Francisco, CA, USA), based on the pTRV2 vector with GenBank accession AF406991. Nuclease-potent *Cas9* line #2 (harboring AtUbqp(1460): human-optimized Cas9/Nosp:NPTII) was used as the platform for generating *eobv*-knockout lines, as described in Skaliter et al. (2024). Sterile explants of *Cas9* line #2 were inoculated with *A. tumefaciens* strain AGL0 harboring pTRV2-*sgRNA1*-s*gRNA2* or pTRV2-*sgRNA1*-*sgRNA3* and pTRV1 (AF406990). Explants were incubated for 20 min in the *Agrobacterium* suspension (in Luria-Bertani broth) containing 100 μm acetosyringone, followed by incubation in the dark for 3 days on Murashige and Skoog (MS) medium supplemented with 3% (w/v) sucrose, followed by transfer for selection on MS with 3% sucrose, antibiotics (kanamycin, carbenicillin) and phytohormones (1.5 mg/L benzylaminopurine [BA], 0.15 mg/L naphthalene acetic acid [NAA]), and incubated under a 12 h light/12 h dark photoperiod at 22°C until regeneration and elongation of regenerants. The medium was renewed every 2 weeks. At the elongation stage, the BA concentration was decreased to 1 mg/L. To switch regenerants to rooting, they were transferred to medium without BA, supplemented with 0.25 mg/L NAA. To screen the gene-edited plantlets, genomic DNA was extracted from them using the CTAB method followed by PCR and Sanger sequencing. Positive plants (T0) were grown on MS with 3% sucrose, transferred to the greenhouse and self-pollinated. Progeny (T1) plants were molecularly analyzed and self-pollinated. The T2 population was screened for homozygous plants, which were selected for further analyses.

#### In silico promoter analysis

Extraction of promoter regions of the *P. axillaris* genome (https://solgenomics.net/) and motif analysis were performed using Python (https://www.python.org/). The code was developed with assistance from ChatGPT and is available at: https://github.com/OdedSkaliter/Promoter-extraction-and-motif-analysis-Petunia-genome.

#### Statistical analysis

Statistical analyses were performed using JMP Pro 18 (SAS, Cary, NC, USA).

## Data availability

The data underlying this article will be shared on reasonable request to the corresponding author.

## Author contributions

O.S. conceptualized, planned and designed the research, performed experiments, analyzed data, and wrote the manuscript. Ek.S., E.L.L., O.R., R.C., El.S., E.M., T.M. and O.E. performed experiments and analyzed data. A.V. conceptualized, planned and designed the research and wrote the manuscript. All authors revised the manuscript and approved the final version.

## Acknowledgments

We thank Danziger – “Dan” Flower Farm for providing petunia cv. Blue Ray plants. We also thank Dr. Guy Polturak (Institute of Plant Sciences and Genetics in Agriculture, The Robert H. Smith Faculty of Agriculture, Food and Environment, The Hebrew University of Jerusalem) for providing *N*. *benthamiana* plantlets and his assistance with the betalain extraction. During the preparation of this manuscript, the authors used ChatGPT (OpenAI, GPT-5.1, 2026) to assist with language editing and text refinement. All AI-assisted content was reviewed and revised by the authors, who take full responsibility for the final version of the manuscript.

## Supplemental material

Figure S1. Virus-induced gene silencing (VIGS) of *AN1* and *AN11* in petunia.

Figure S2. Yeast-two hybrid assay testing the interactions of various MYB regulators of volatile production with AN1 and AN11.

Figure S3. Protein domain analysis of EOBV.

Figure S4. EOBV participates in the MBW complex.

Figure S5. EOBV is an R2R3-MYB subgroup 4 member.

Figure S6. *EOBV* expression levels in petunia floral buds and flowers 1 day postanthesis (1DPA) in the morning (1000 h) and evening (1900 h).

Figure S7. Virus-induced gene silencing (VIGS) of *EOBV* in petunia flowers.

Figure S8. Effect of exogenous gibberellin (GA) application on *EOBV* transcripts.

Figure S9. In silico promoter analysis in the *Petunia axillaris* genome.

Figure S10. Diel, developmental and spatial expression patterns of PhMYB4.

Figure S11. *PhMYB4* genomic sequence is not affected by knockout of *eobv*.

Figure S12. Stacking of *ever-* and *eobv*-knockout alleles.

Figure S13. Effect of elevated growth temperature regime on expression of *PhMYB4* and α-expansin (*PhEXP1*) in *eobv*-knockout lines.

Supplementary Table 1. Promoters of genes in the *Petunia axillaris* genome containing the motif YACCWACY.

Supplementary Table 2. List of primers used in this study.

