## Supplementary material for "The MBW complex regulates volatiles in petunia flowers: EOBV interacts with AN1 to suppress biosynthesis of phenylpropenes": Supplementary Figures.pdf

**Running title: EOBV is a component of the MBW complex regulating scent**

a

*AN1*:
GGTAGAGAGGTGTGATTGTGGGGAACAACTCCTGTATTAACGGT
CTTTTCTGATATACAGGTGACTTGTCTAACGTGTCGCAAGGCATT
GTACCAAACGTCGACGTAATAATAAACATGACGTTGGTCTTTGAC
GTGCTATTATAACCTTTCCTCTAATGAAATAAGATGTGGATtATTAAT
ATCACTGATTCTTTTTATTTGTATAATCAACTAGTTGTTTAC

b

*AN11*:
TATGCTCTTTTGCCACACAGAATAAGTTGTAACTATCTAGTAGTAGCAGCAGAAGCAAT
AGAACCTATCTTACATGAAAAAGATTCAATCTTTCAATCACCTTATACTTTAAGCAATTG
CAACTTGTTAGAAAATGCAATAGCAATCCAATCACGCTGCGCAGGAGACCACTGAAGC
TGATTAATCTCAGCACCAGCAGAGTACATCGACATGGGATCAATCCCATTAGGCCCAG
CAACAGTAGGCAACTCCCAAATCAACGCCTGCCCATCATCTCCACCTGAACAAATATG
TCTACAACTCTGGGGAGCCCAAGCAATGGCATTCACACT

c

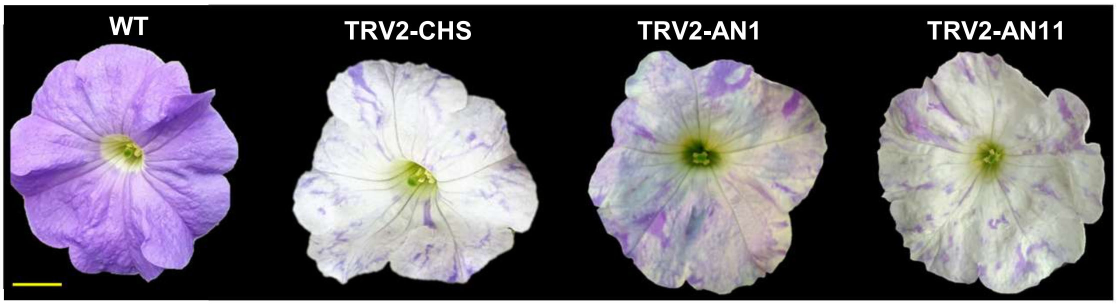


**Figure S1. Virus-induced gene silencing (VIGS) of *AN1* and *AN11* in petunia. (a and b)** Sequences used for VIGS of 5' untranslated region (UTR) of *AN1* **(a)** and 3' UTR of *AN11* **(b)**. **(c)** Representative photos of wild-type (WT) petunia cv. P720 flower, and corollas displaying white sectors due to loss of anthocyanins as a result of *CHS*, *AN1* or *AN11* suppression. Bar = 1 cm.

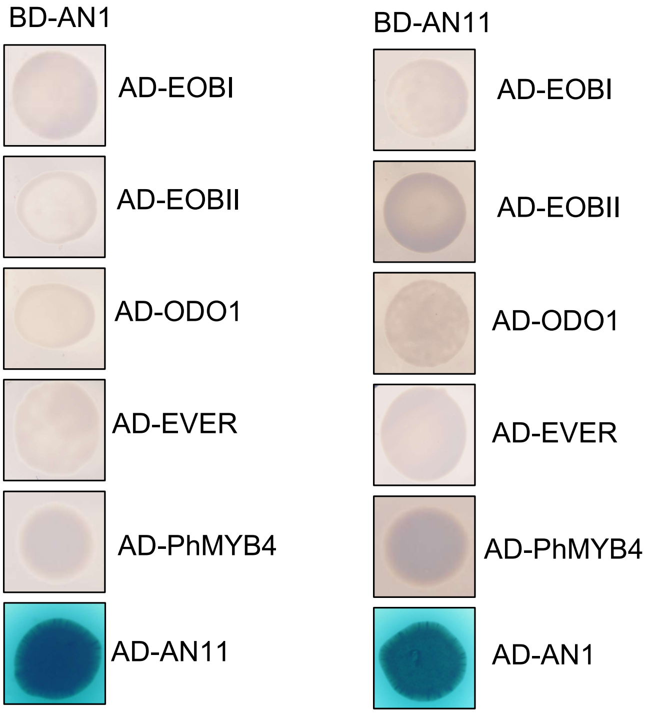


**Figure S2. Yeast-two hybrid assay testing the interactions of various MYB regulators of volatile production with AN1 and AN11.**

The coding sequences of the targets were fused to GAL4 binding domain (BD) or GAL4 activation domain (AD) and transformed into yeast. X-Gal staining was performed on transformed yeast to detect $\beta$-galactosidase activity, indicating an interaction between bait and prey. The interaction between AN1 and AN11 was used as a positive control. EOBI, EMISSION OF BENZENOIDS I; EOBII, EMISSION OF BENZENOIDS II; ODO1, ODORANT1; EVER, EPIDERMIS VOLATILE EMISSION REGULATOR.

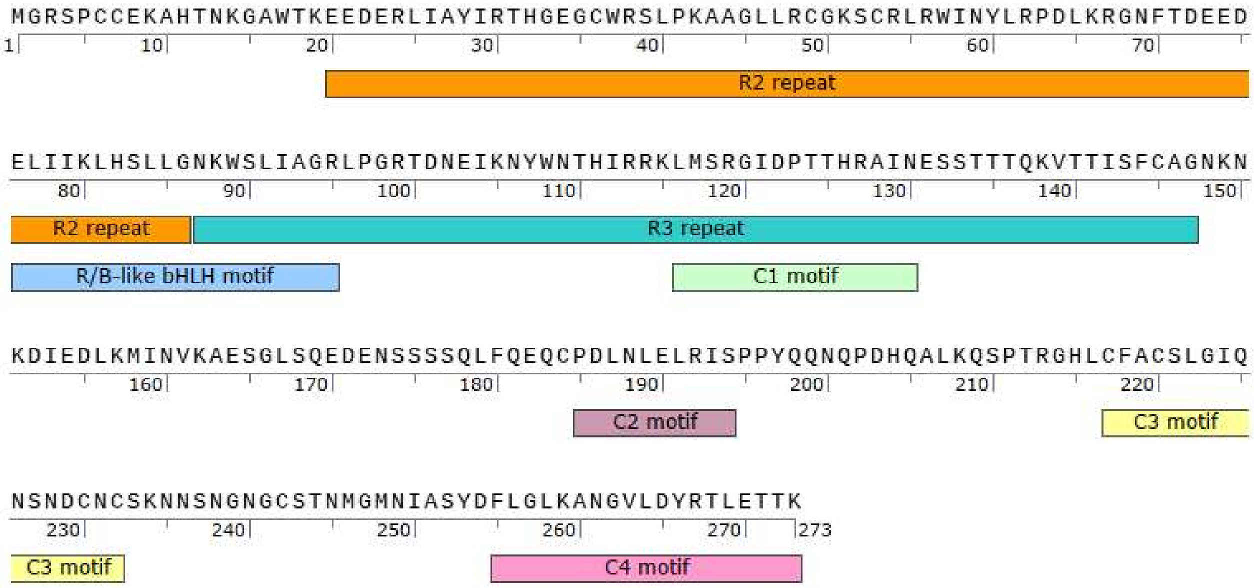


**Figure S3. Protein domain analysis of EOBV.**
The analysis was performed using InterPro (Blum et al., 2021). Domains are color-keyed. R/B-like bHLH motif – D/E]Lx2[R/K]x3Lx6Lx3R; C1 motif – LlsrGIDPxT/sHRxI/L; C2 motif – pdLNLD/ELxiG/S; C3 motif – CX1-2CX7-12CX2C; C4 motif – FLGLx4-7V/LLD/GF/YR/Sx1LEMK.

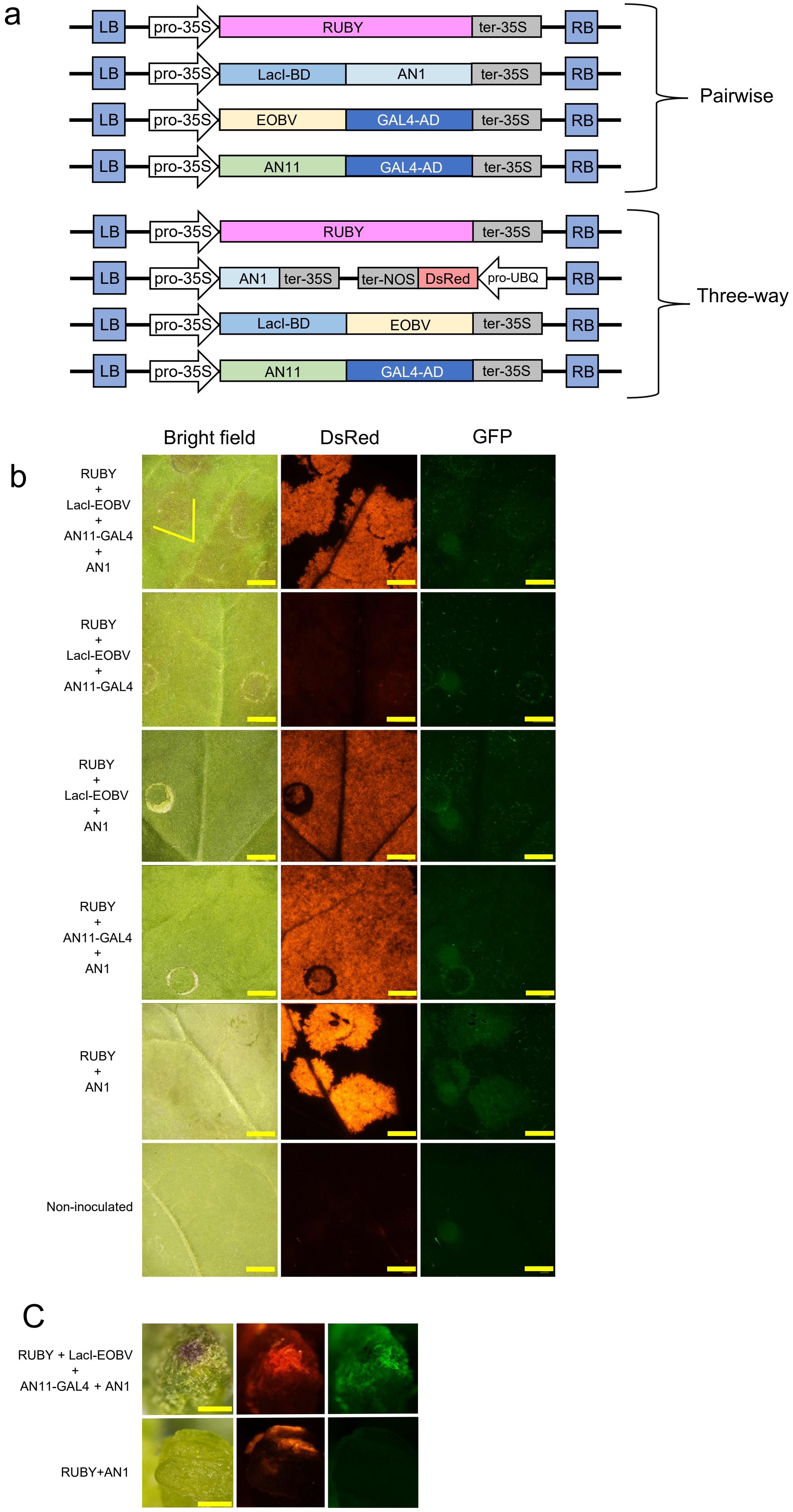
a
LB
pro-35S
RUBY
ter-35S
RB
LacI-BD
AN1
EOBV
GAL4-AD
AN11
Pairwise
ter-NOS
DsRed
pro-UBQ
Three-way
Bright field
DsRed
GFP
b
RUBY + LacI-EOBV + AN11-GAL4 + AN1
RUBY + LacI-EOBV + AN11-GAL4
RUBY + LacI-EOBV + AN1
RUBY + AN11-GAL4 + AN1
RUBY + AN1
Non-inoculated
C
RUBY + LacI-EOBV + AN11-GAL4 + AN1
RUBY+AN1

**Figure S4. EOBV participates in the MBW complex.**

**a)** Schematic representation of vectors used in the protein–protein interaction assays. Top: vectors used in pairwise interaction assays. Bottom: vectors used in three-way interaction assays. OpLacI:mini35S, 6 repeats of the lac operator fused to a minimal 35S promoter. The coding sequences of the targets were fused to LacI binding domain (BD) or GAL4 activation domain (AD). **b)** *Nicotiana benthamiana* leaves were inoculated with a mixture of agrobacteria carrying different vector combinations; 3 days postinoculation, leaves were examined under a fluorescent stereomicroscope. Yellow arrows mark observable betalain accumulation in leaves inoculated with OpLacI:RUBY, 35S:LacI-EOBV, 35S:AN11-GAL4 and 35S:AN1. Bar = 1 mm. **c)** Representative photos of pigmented regions formed on *Nicotiana tabacum* leaf disks inoculated with OpLacI:RUBY, 35S:LacI-EOBV, 35S:AN11-GAL4 and 35S:AN1 and control inoculated with OpLacI:RUBY and 35S:AN1. Bar = 1 mm.

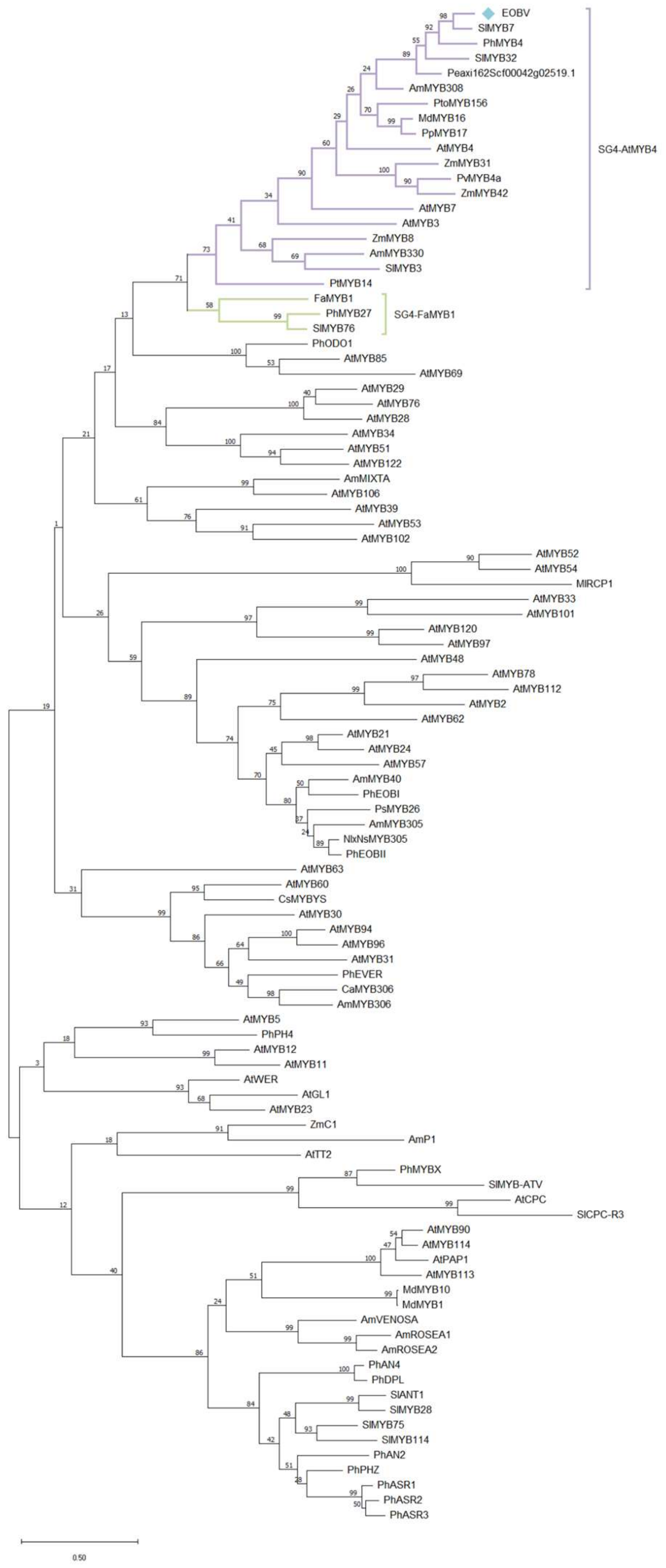


**Figure S5. EOBV is an R2R3-MYB subgroup 4 member.**

Phylogenetic analysis representing the relationships between EOBV (marked by a diamond) and R2R3-MYBs from petunia and other plant species. Subgrouping is based on Dubos et al. (2010). Protein sequence alignment was done by ClustalW followed by maximum-likelihood method with 1000 bootstraps using MEGA version 11. Bootstrap values are indicated at the branch nodes, and scale bar indicates the number of amino acid substitutions per site.

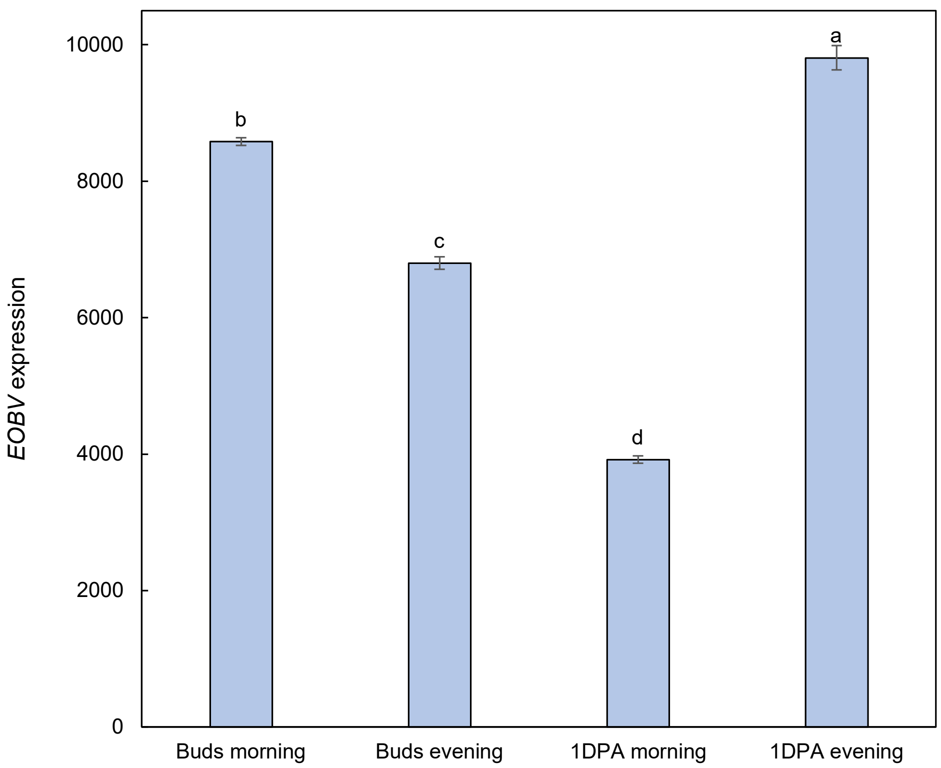


**Figure S6.** ***EOBV*** **expression levels in petunia floral buds and flowers 1 day postanthesis (1DPA) in the morning (1000 h) and evening (1900 h).**

Transcripts were detected by RNA-sequencing performed in Shor et al. (2023b). Data are means ± SEM ($n$ = 3). Significance of differences was calculated by Tukey's multiple comparison test following one-way ANOVA. Values with different letters are significantly different at $P \leq 0.05$. Standard errors are indicated by vertical lines.

a

AAAACAGAAACCAAAGAACAGAGAGGAAGAAAAGAATATACAAGGAATAGGATCAT
GAGTCGAGGTATTGACCCTACAACACACAGGGCTATTAACGAGTCTAGTACTACCACAC
AAAAAGTTACAACAATTTCTTTTTGTGCTGGAAATAAAAATAAAGATATTGAAGATCTG
AAGATGATCAATGTCAAAGCTGAATCTGGACTTAGCCAAGAAGATGAAAATAGTAGCA
GCAGCCAATTATTTCAAGAACAGTGTCCTGATTTGAATCTTGAGCTCAGAATTAGCCCT
CCTTACCAACAAAACCAACCAGATCACCAAGCTTTGAAACAAAGTCCAACAAGGGGC
CATTTGTGTTTTGCATGCAGTTTGGGTATCCAAAACAGTAACGATTGCAATTGCAGTAA
AAATAATAGTAATGGGAATGGTTGCAGTACTAACATGGGTATGAATATTGCAAGTTATG
ATTTTTTAGGATTAAAAGCTAATGGTGTTTTGGACTACAGAACCTTGGAGACTACTAAG
TGATTATTATGTTGGATTACATAAAAAAAAAGAGAATTGAGAAAGACAGTGTTATTAAT
TTAAGTCTTTTTCGAATTTCTCCTATTTGTAAAGTTGAAAGTAATATATAGTAGTATTACA
TTAAGTTGAAAGCAGTACATCAGTGTACCTTATACTAAATTTTAATT

b

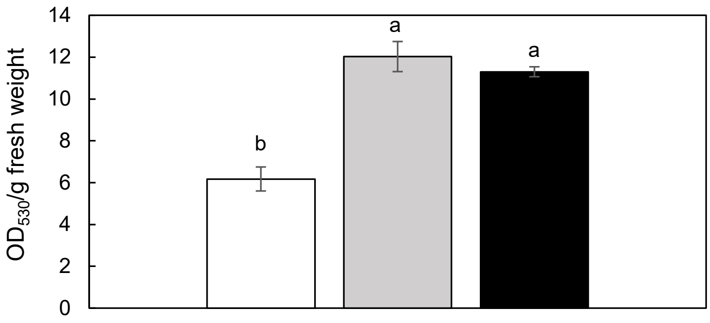


**Figure S7. Virus-induced gene silencing (VIGS) of *EOBV* in petunia.**
**a)** mRNA sequence of *EOBV*. Red nucleotides, 5’ untranslated region (UTR); black nucleotides, coding sequence; green nucleotides, 3’ UTR used for VIGS.
**b)** Anthocyanin content in corollas of petunia cv. Classic Blue Ray accumulating purple anthocyanins following suppression of *EOBV*. Plants (1 month old) were inoculated with agrobacteria carrying pTRV2-CHS as a control or pTRV2-EOBV. After 1 month, white petal tissue (indicating *CHS* silencing, white bar), non-suppressed purple petal tissue (grey bar) and petal tissue of plants inoculated with TRV2-EOBV (black bar) was collected, and anthocyanins were extracted and analyzed spectrophotometrically. Data are means ± SEM ($n$ = 4). Significance of differences was calculated by Tukey's multiple comparison test following one-way ANOVA. Values with different letters are significantly different at $P \leq 0.05$. Standard errors are indicated by vertical lines.

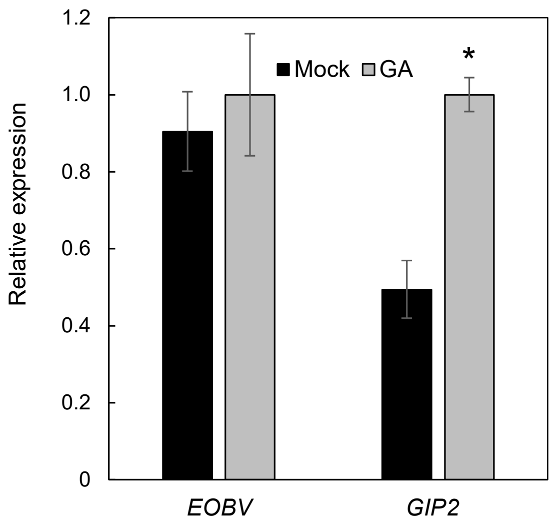


**Figure S8. Effect of exogenous gibberellin (GA) application on *EOBV* transcripts.**

Petunia buds were treated with or without (mock) $GA_3$ and after 48 h, RNA was extracted from the flowers and subjected to RT-qPCR analyses. Data were normalized to *UBIQUITIN* and to the sample with the highest expression level. Data are means ± SEM ($n$ = 3). Significance of differences between treatments was calculated by two-tailed unpaired Student's t-test: *$P \leq 0.05$. Standard errors are indicated by vertical lines. GIP, GIBBERELLIN-INDUCED PROTEIN 2.

a

*C4H2* (Peaxi162Scf00390g00225):
ACAATATCCAACTCATAATCTTATCAATCTTGTTTGAGTAATCAACATTAATT
CTTATATGCATCTAGTAAGTTCTTTTACAAAAACT**CACCAACC**TTTCCACT
ATTTCTTTTTCCTCTACCAAACCTCCAATCTACCATCTTATCAACTCCGTC
AAACTTAACACCGTTAACCTCCC**CACCTACC**CCCACCAGTTTCCCTCTAT
ATAAATCCACATATACACACCTTTATCAGCCGTACCAAACAAACTCACTTT
TATTGTCCAAACGAAAAAAAAAAAAAGAACTTGTCCAAACAAAA

*PAAS* (Peaxi162Scf00561g00021):
AAGTAAGGCTAGTATTCTTTTGTGTAAATCAATCTACCTTACTCATCCACC
AAGGTTGAGATAGTGTGGAATCGGATTTAATAAATATCTATGTCATTTTTTC
ACATAGAGGTCTTGCAAATATTTTCAGTACAATTCAATTTTATCATAAGTTA
CCAAAGGTGTATAGCTGGATCTATATATAATTAGATTGCTAAGTTCATGTTG
CACATTCATATTATAATACTCTCTTTGTTTGTTCCCTTCTCTGCTTTTAGTTT
AATTGTACGTCATATACCTTCAAATTCAAAATTTATTTCC

*BSMT* (Peaxi162Scf00047g01123):
TTACTCCAATACCCGTTCGAATTTTGAATCTCCAAAATAAAAGAATTGACT
TCAAATGAGAACACATAAGAATTGAAACAATAACTAAAGCCTCTTACTAAG
TTTATAATATAGATTTAGACCACTTACTCATCTATATCCCTGTCACTGTCAG
GGCCAATACTGGAATTTTAGTGGACAAACTTCAGCAGTAGCACCTTTCTA
TAAATAGGTAATAAATTCTTGCCATGTAGAAGCACTATCTCACATAAAGAAT
AACAAACAAAAAGATCAAGAAAGAGATACTCATAGCAAGAAGAA

b

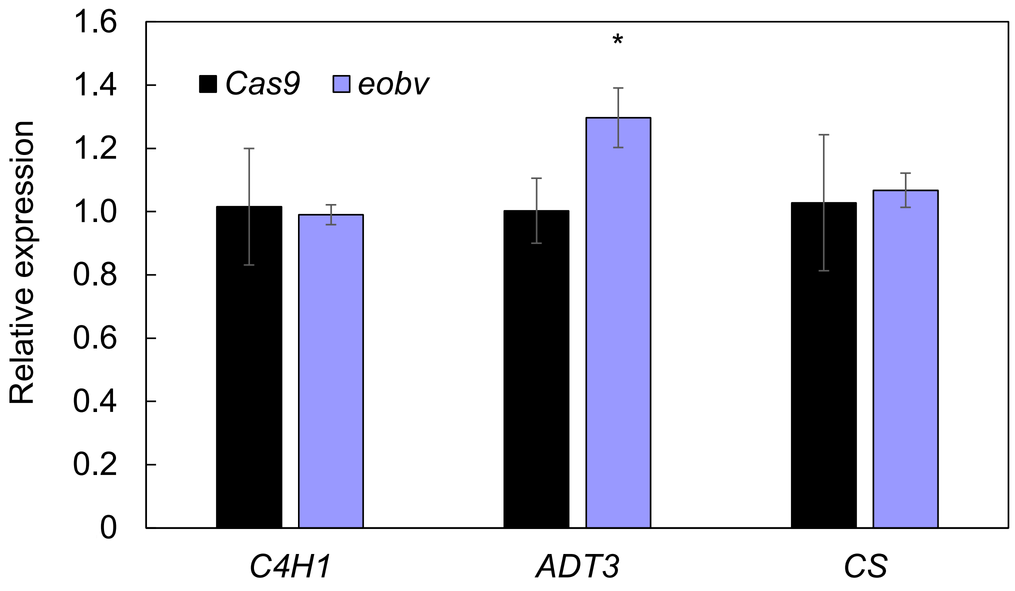


**Figure S9. In silico promoter analysis in the *Petunia axillaris* genome.**
**a)** Promoter sequences of *C4H2*, *BSMT* and *PAAS*. Nucleotides marked in red indicate the conserved motif YACCWACY identified by AtMYB4. Sequences were obtained from Sol Genomics Network (https://solgenomics.net/). **b)** Expression of selected genes containing the YACCWACY motif, as identified in the genome-wide analysis, in *eobv*-knockout lines vs. *Cas9* controls**.** RT-qPCR analysis was performed on RNA extracted from flowers at 2 days postanthesis. Data are means ± SEM ($n$ = 3). Data were normalized to the geomean of *EF1α* and *UBIQUITIN*. Standard errors are indicated by vertical lines. Significance of differences between treatments was calculated by two-tailed unpaired Student's t-test: *$P \leq 0.05$. C4H, CINNAMATE 4-HYDROXYLASE; PAAS, PHENYLACETALDEHYDE SYNTHASE; BSMT, S-ADENOSYL-L-METHIONINE:BENZOIC ACID/SALICYLIC ACID CARBOXYL METHYLTRANSFERASE; ADT, AROGENATE DEHYDRATASE; CS, CHORISMATE SYNTHASE.

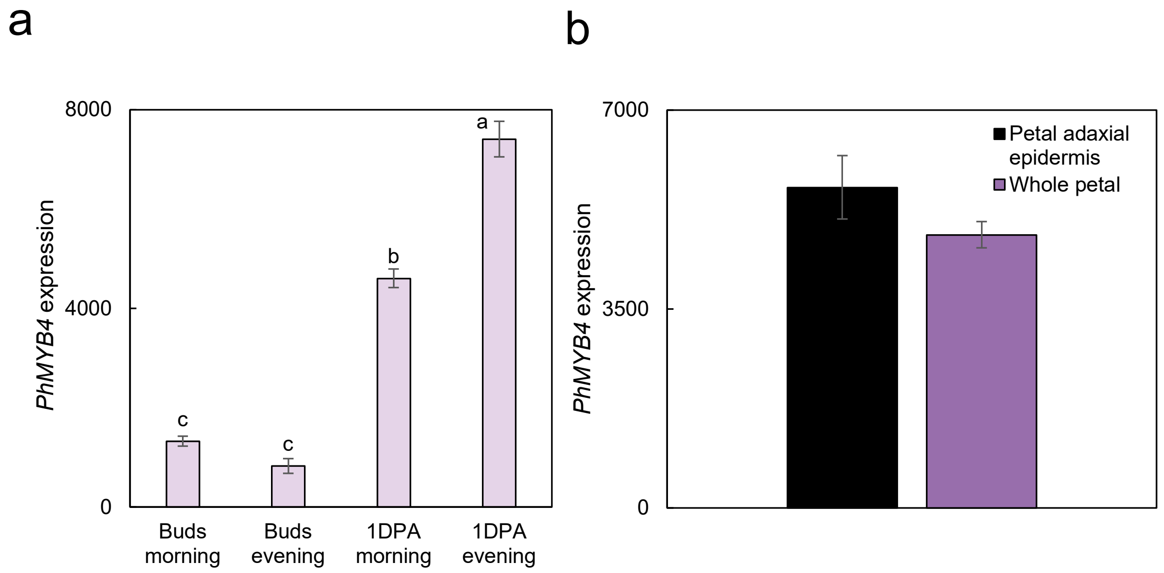


**Figure S10. Diel, developmental and spatial expression patterns of *PhMYB4*.**

**a)** *PhMYB4* expression levels in petunia floral buds and flowers 1 day postanthesis (1DPA) in the morning (1000 h) and evening (1900 h). Transcripts were detected by RNA-sequencing performed in Shor et al. (2023b). Significance of differences was calculated by Tukey's multiple comparison test following one-way ANOVA. Values with different letters are significantly different at $P \leq 0.05$.

**b)** *PhMYB4* transcript levels in the petal adaxial epidermis. Normalized counts in the petal adaxial epidermis from the RNA sequencing performed in Skaliter et al. (2024). Data are means ± SEM ($n$ = 3). Standard errors are indicated by vertical lines.

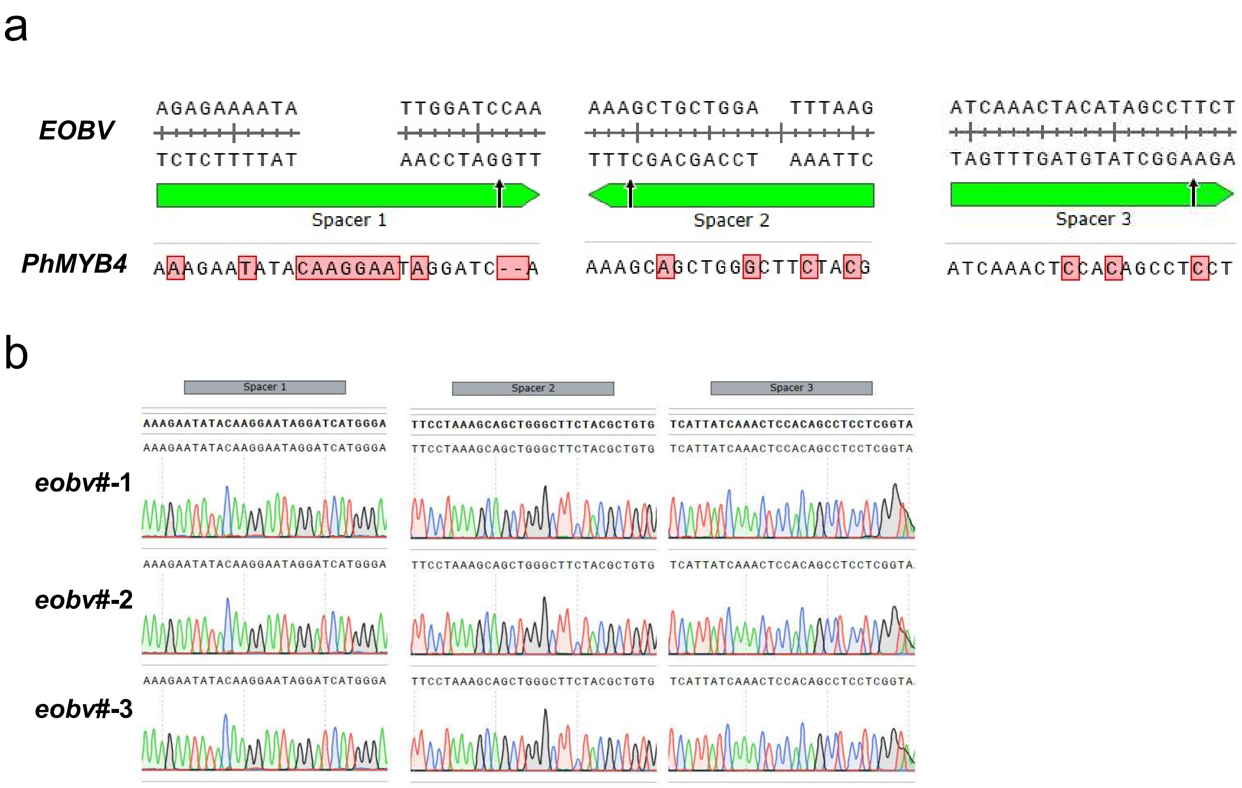


**Figure S11. *PhMYB4* genomic sequence is not affected by knockout of *eobv*.**

**a)** Alignment of spacer sequences designed to target *EOBV* and *PhMYB4* genomic sequence.

**b)** Sanger sequencing of *PhMYB4* in *eobv* knockout lines #1–3 showing no editing events. Gray rectangles mark the location of the *PhMYB4* sequences with the highest homology to the spacers designed to target *EOBV*.

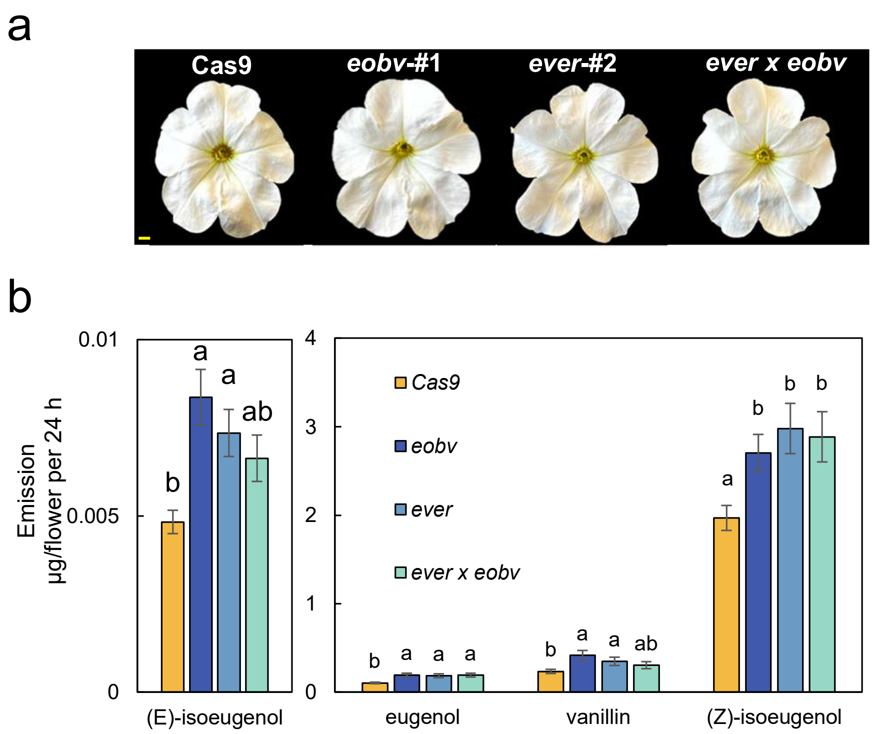


**Figure S12. Stacking of *ever* and *eobv* knockout alleles. a)** Representative petunia cv. Mitchell flowers from *Cas9* control, *eobv*-#1, ever-#2 and *ever* x *eobv* at 2 days post anthesis (2DPA). Bar = 1 cm. **b)** Dynamic headspace (collected for 24h and analyzed by GC-MS) was performed on flowers at 2DPA collected from *Cas9* (control), *ever*-#2, *eobv*-#1 and *ever* x *eobv.* Data are means ± SEM (*n* = 15-17). Significance of differences was calculated by Tukey's multiple comparison test following one way analysis of variance. Values with different letters are significantly different at $P \leq 0.05$. Standard errors are indicated by vertical lines.

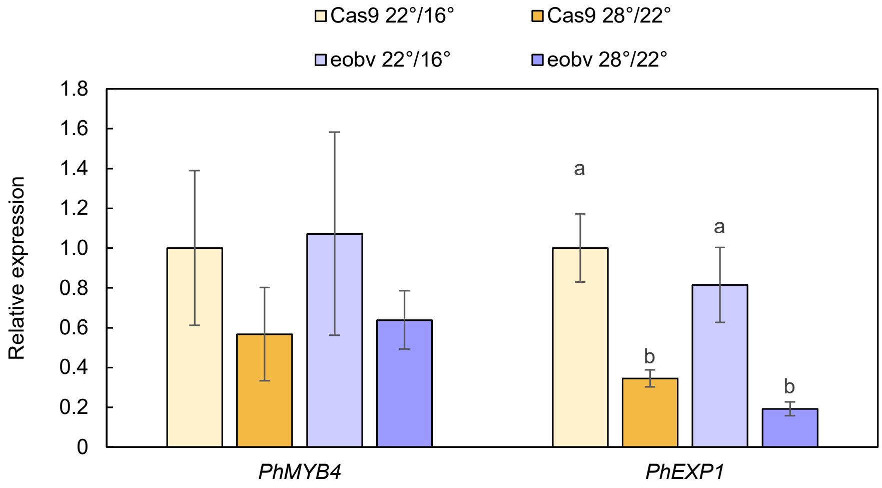


**Figure S13. Effect of elevated growth temperature regime on expression of *PhMYB4* and *α-expansin* (*PhEXP1*) in *eobv*-knockout lines.**

Plants of *eobv*-knockout lines and control Cas9 were grown for 1 month in a phytotron at different growth temperatures: 22/16°C and 28/22°C. RT-qPCR analysis was performed on RNA extracted from flowers at 2 days postanthesis. Data are means ± SEM ($n$ = 4–5). Data were normalized to the geomean of *EF1α* and *UBIQUITIN*. Standard errors are indicated by vertical lines. Significance of differences was calculated by Tukey's multiple comparison test following one-way analysis of variance. Values with different letters are significantly different at $P \leq 0.05$.
